# Genetic diversity, population structure and differentiation of farmed and wild African catfish (*Clarias gariepinus*) in Nigeria

**DOI:** 10.1101/2024.10.15.618429

**Authors:** Mark K. Sanda, Neil B. Metcalfe, Maria Capstick, Jenna Nichols, Barbara K. Mable

## Abstract

The African catfish (*Clarias gariepinus*) is a commercially important species, for both fisheries and aquaculture, and is now the most commonly farmed fish in sub-Saharan Africa. However, knowledge about the genetic diversity and population structure of wild and farmed populations, which is crucial for effective conservation and sustainable aquaculture management, is scarce. Using mitochondrial DNA (mtDNA) cytochrome c oxidase 1 gene (COI) sequencing and genomic analysis using triple restriction site-associated DNA sequencing (3RAD), we investigated the genetic diversity and population structure of farmed and wild *C. gariepinus* populations from Nigeria, including an albino form found in the wild. Eleven COI haplotypes were identified, of which seven were unique to wild samples. Wild sampling sites had a slightly broader range and higher maximum values for observed heterozygosity (*Ho* = 0.109 – 0.165), expected heterozygosity (*He* = 0.111 – 0.216), and nucleotide diversity (*pi* = 0.125 – 0.225) compared to the farmed populations (*Ho* = 0.118 – 0.147, *He* = 0.112-0.144, *pi* = 0.117 – 0.151). Conversely, genetic differentiation (*Fst*) was higher among farmed sampling sites compared to the wild ones and there was high genetic differentiation between the farmed and wild *C. gariepinus* sampling sites (*Fst* = 0.31 – 0.47). Despite evidence for admixture for both farmed and wild fish, there was little evidence of admixture between the two groups. Nevertheless, both mtDNA and 3RAD data strongly suggested that the albino fish, collected from the wild, were in fact farm escapees. Despite overall differentiation farmed genotypes suggesting that overall genetic integrity of wild fish has been maintained, this evidence of escape provides a warning about potential risks of increasing aquaculture activities. Specifically, this indicates the need for greater regulation of fish farms to monitor and reduce the risk of escapes.

## 1 Introduction

The African catfish, also known as the African sharptooth catfish or the North African catfish, is a freshwater omnivorous species that belongs to the Claridae family and is found in tropical and subtropical climates (Omitoyin, 2007). Its native distribution range spans lakes, reservoirs, and rivers in sub-Saharan Africa and it also has been introduced into South America, Southeast Asia, and Europe (Konings et al., 2019; Truter et al., 2023). It is an important commercial fish both as a capture and aquaculture species, and is now the most farmed species in sub-Saharan Africa (Chandra Segaran et al., 2023). *C. gariepinus* has drawn the attention of aquaculturists because of its biological attributes, which include a fast growth rate, resistance to diseases and tolerance of high stocking densities (Lal et al., 2003). Research on *C. gariepinus* has led to the development of a strain known as the “Dutch *Clarias*” through selective breeding conducted in Belgium and the Netherlands, following their introduction to those countries from Africa. Dutch *Clarias* have since been introduced to different African countries including the Central African Republic, South Africa, Côte d’Ivoire, and Nigeria, where they are cultivated as farmed fish for food (Holčík, 1991; Huisman and Richter, 1987; Roodt-Wilding et al., 2010).

There is a tension between the use of *C. gariepinus* in aquaculture and its conservation in the wild, since it is essential to maintain a balance between exploiting its economic potential and preserving the genetic diversity of wild stocks. Previous studies have revealed that wild *C. gariepinus* populations exhibit high genetic variation, which is an important parameter for the species’ adaptability and resilience to changing environmental conditions (Barasa et al., 2014; Barasa et al., 2017). Given the importance of genetic diversity as a valuable resource for fish species, it is therefore the responsibility of policy makers and managers to define fisheries and aquaculture programmes that will ensure the sustainable utilisation of economically important species like *C. gariepinus*. Conservation officers must therefore learn to deal with factors posing threats to the genetic conservation of natural resources including loss of genetic diversity due to hybridisation with other species, as observed in Bangladesh with *C. gariepinus* and *C. batrachus* (Parvez et al., 2022), habitat fragmentation, overfishing, and pollution. Additionally, the use of Dutch Clarias in fish farms outside their native range could negatively impact biodiversity in the event of fish escapes, as observed in Brazil and Turkey (Dumith and Santos, 2022; Turan and Turan, 2016).

*C. gariepinus* is by far the most commonly farmed fish in Nigeria (FAO, 2022). Both wild-caught *C. gariepinus* and the introduced Dutch *Clarias* are widely used in artificial breeding programmes, with reproduction being induced through hormone treatment with Ovaprim (Ataguba et al., 2009). *Clarias gariepinus* has been identified as a cheap source of animal protein and means to achieving food security, as well as job creation (Folorunso et al., 2021). Given their high economic importance to the country, it is important that the conservation of *C. gariepinus* and their sustainable exploitation in the wild is prioritised. However, there is limited knowledge on the genetic diversity of either wild or farmed catfish populations in the country. This is coming at a time when fish production in Nigeria is facing threats from environmental change (evident in the receding of freshwater lakes and rivers due to extended dry seasons), overexploitation, and lack of enforcement of the policy guiding fishing activities (Olopade et al., 2017). As a result, wild fish harvests in Nigeria have been declining in recent years (FAO, 2022). The factors linked to the reduction in fishing yields in Nigeria have elsewhere been reported to have negative effects on the genetic diversity of wild fish populations (Coleman et al., 2018; Kenchington, 2003; Yan et al., 2021). Genetic diversity is crucial for the long-term survival, adaptation, and resilience of individuals, populations, species, and ecosystems, as it forms the foundation of biodiversity (Hvilsom et al., 2022). Past studies conducted on *C. gariepinus* in Nigeria have attempted to identify and assess the genetic diversity of *C. gariepinus* species using DNA markers. For example, Suleiman et al. (2020) observed high genetic diversity within farmed and wild *C. gariepinus* populations from the northeast of Nigeria using random amplified polymorphic markers (RAPD). Popoola (2022) observed high genetic differentiation among three wild *C. gariepinus* populations from Southwestern Nigeria using sequences from the mitochondrial cytochrome b (cytb) gene. This molecular approach has been extended to identify wild freshwater fish in Southeastern Nigeria using the mtDNA marker cytochrome c oxidase subunit I (COI) (Nwani et al., 2011). Awodiran et al. (2019) provided useful information on genetic diversity for management and conservation of fish while attempting to differentiate between wild *C. gariepinus* populations from Northcentral and Southwestern Nigeria using microsatellite markers. These past studies have offered information on native species diversity but with limited geographical coverage; moreover, they did not compare farmed and wild populations to assess the levels of genetic differentiation and hybridisation, or detection of possible escapes from farmed to wild.

These issues can now be tackled in greater depth due to technological advances. For example, the use of high-throughput sequencing technologies and genomic approaches in fisheries and aquaculture provide a means of identifying population structure, genetic diversity and differentiation among different fish populations, which can be used to inform fisheries management and identify escapees from fish farms (Bernatchez et al., 2017). The advancement in next-generation sequencing has led to the assembly of over 270 fish genomes to promote studies on comparative genomics, evolution, and systematics and more importantly for its application in aquaculture and fisheries (Bian et al., 2019; Crollius and Weissenbach, 2005; Hughes et al., 2018; MacKenzie and Jentoft, 2016). However, despite its relevance as one of the most important farmed species in sub-Saharan Africa (Hecht, 2013; Mramba and Kahindi, 2023), and decades of domestication (Hecht, 2013), it was only in 2022 that the Leibniz Institute for Farm Animal Biology (FBN) sequenced and assembled the genome of *C. gariepinus* (GCA_024256425.2).

Whole genome sequencing provides comprehensive information on genetic variation, including single nucleotide polymorphisms (SNPs), insertions, deletions, and structural variants (Ng and Kirkness, 2010). Due to the high cost of sequencing the entire genome of an organism, alternative approaches like reduced representation sequencing technologies, based on high-throughput SNP genotyping from DNA fragmented with restriction enzyme(s), have made it possible to study a fraction of genomes (Peterson et al., 2012). The most employed technique for reduced-representation sequencing is known as restriction site-associated DNA sequencing (RADseq). Several RADseq methods have evolved over the years, including RADseq based on one restriction enzyme (Baird et al., 2008), double digest restriction site-associated DNA (ddRAD) sequencing (Peterson et al., 2012) and most recently, triple-enzyme restriction site-associated DNA sequencing (3RAD) (Bayona-Vásquez et al., 2019). The 3RAD approach is a low-cost, highly robust and simple method for the construction of dual-digest RADseq libraries using 96 pairs of Illumina compatible iTru5 and iTru7 primers enabling multiplexing of more samples and pooling of more libraries than other RADseq approaches (Bayona-Vásquez et al., 2019). The 3RAD approach enables the simultaneous digestion and ligation of DNA, which minimises library construction steps. Integrating the mtDNA COI marker with the cost-effective 3RAD genomic approach offers a promising strategy to improve the accuracy and robustness of population genetic analyses in *C. gariepinus*.

Here, we aimed to compare the genetic diversity and differentiation among farmed and wild *Clarias gariepinus* populations in Nigeria, using a combination of mtDNA barcoding with the COI marker and 3RAD high-throughput sequencing. Specifically, we assessed whether: 1) mtDNA haplotype distributions in Nigeria, in combination with published information in the Barcode of Life Database (BOLD) based on 100% sequence match, were informative about the genetic integrity of wild fish; 2) there was evidence of differentiation between farmed and wild fish based on mtDNA; 3) genome-wide patterns of polymorphism based on 3RAD analyses indicated differences in levels of genetic diversity or inbreeding in farmed vs wild fish; and 4) there was evidence of genetic differentiation or admixture among wild and farmed sites based on genome-wide SNPs.

## 2 Methods

### 2.1 Sample collection

Caudal fin clip samples of *C. gariepinus* were collected from northeastern and southwestern regions in Nigeria during a four-month period between November 2021 and March 2022 (Figure 1, Table 1). The sampling was conducted in four farmed sites (southwest: f_CMC, f_ODC, f_LAC; northeast: f_SAC) and five wild sites in the northeast with three sites from Adamawa state (w_BYC: River Benue Yola, w_KDC: Kiri dam, and w_LGC: Lake Geriyo) and two sites from Gombe state (w_DKAL: albino samples from Dadin Kowa dam, w_DKC: Dadin Kowa dam non-albino samples). Lake Geriyo is a tributary to River Benue while Dadin Kowa dam is located on the Gongola River, which is a major tributary of the Benue River (Essien et al., 2019; Hassan et al., 2015). A total of 222 fin clips were preserved in RNAlater (Invitrogen, United Kingdom) and stored in a refrigerator before DNA extraction. Samples for genetic analysis were chosen by aiming for a subset of 15 randomly-selected fish from each sampling site. However, the actual sample sizes varied among sites due to DNA degradation issues. For sites with sample sizes less than 15, such as the albino (n = 5) and River Benue Yola (n = 9) sampling sites, we took genetic samples from all available fish to maximise the representation of genetic diversity within these sites. This approach allowed us to adapt to the observed sample sizes and DNA quality constraints while ensuring adequate representation for downstream genetic analysis.

**Figure 1.**
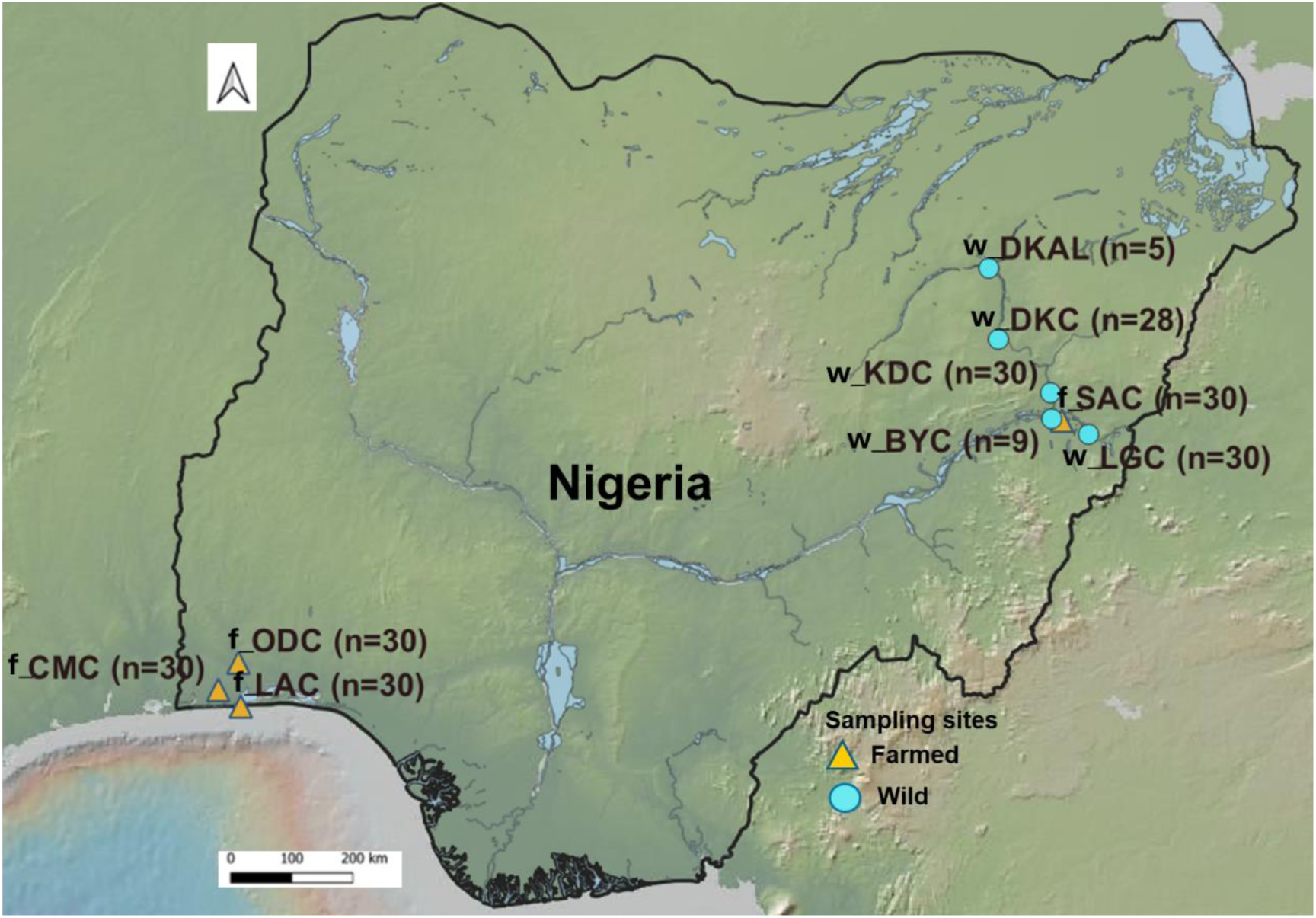
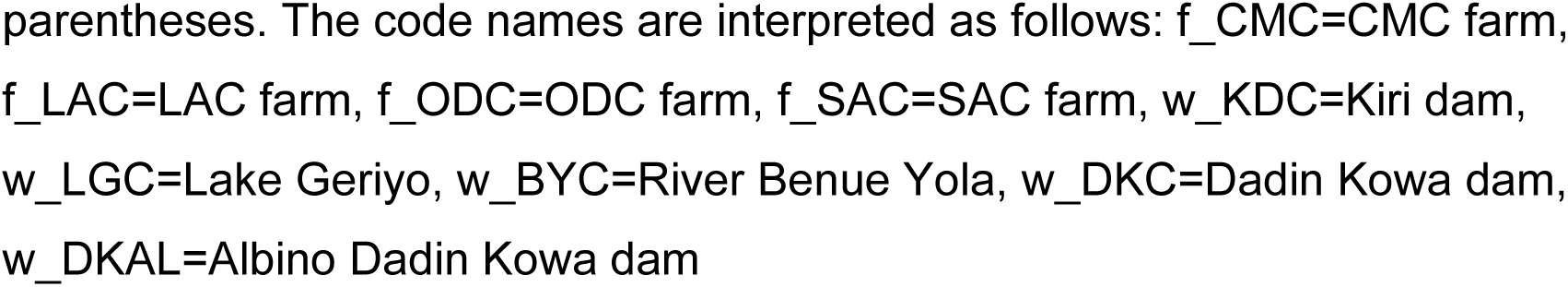
Map of Nigeria illustrating the spatial distribution of water bodies and sampling locations within the study area, categorised based on the origin of the sampled *Clarias gariepinus* populations (farmed and wild), with the sample size in parentheses. The code names are interpreted as follows: f_CMC=CMC farm, f_LAC=LAC farm, f_ODC=ODC farm, f_SAC=SAC farm, w_KDC=Kiri dam, w_LGC=Lake Geriyo, w_BYC=River Benue Yola, w_DKC=Dadin Kowa dam, w_DKAL=Albino Dadin Kowa dam

**Table 1.**
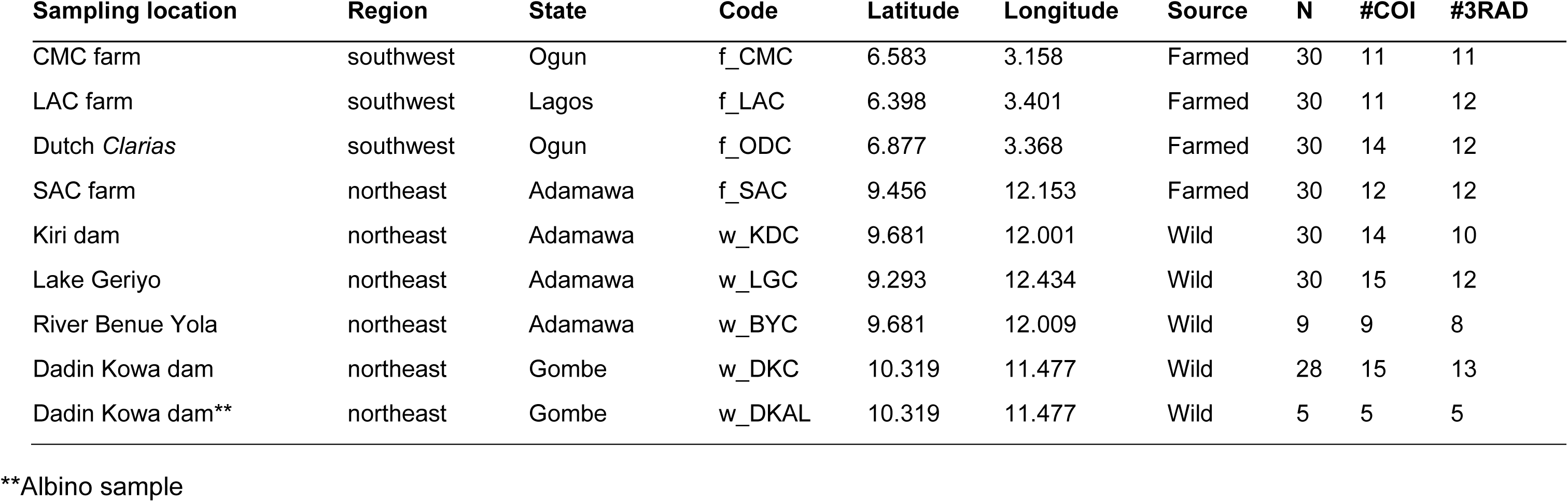
Origin of farmed and wild *Clarias gariepinus* collected from various states in Nigeria, indicating the sampling locations, region, state, sampling site code, geographical coordinates, whether the population is farmed or wild, the total sample size collected per population (N), and sample sizes for mitochondria DNA barcoding (#COI) and 3RAD (#3RAD) analyses. The Dadin Kowa dam with an asterisk are albino samples, which were considered separately from the other samples in the population.

### 2.2 DNA isolation and PCR amplification

Genomic DNA was extracted using DNeasy Blood & Tissue Kits (Qiagen Inc, Paisley, UK), following the manufacturer’s instructions for tissue samples, with elution into 100 µl of AE buffer. The integrity of the extracted DNA was verified in 2% agarose gel electrophoresis and the concentrations (ng/µl) were measured using a Qubit 2.0 Fluorometer (Thermo Fisher Scientific, Waltham, Massachusetts, USA), with the broad-range kit (Invitrogen, Massachusetts, USA). The cytochrome oxidase I (COI) gene was amplified using the primer pair FISH-BCL 5’-TCAACYAATCAYAAAGATATYGGCAC-3’ and FISH-BCH 5’-TAAACTTCAGGGTGACCAAAAAATCA-3’ (Baldwin et al., 2009). A 20 μL PCR master mix was prepared containing 15 μL ddH_2_O, 2 μL 10X buffer, 1 μL 50mM MgCl_2_, 0.4 μL 10mM dNTPs, 0.2 μL 10mM primer F, 0.2 μL 10mM primer R, 0.2 μL of 5,000 units/ml Taq (Invitrogen, United Kingdom), and 1 μL DNA. PCR was performed under the following conditions: initial denaturing at 94°C for 4 min, 52°C for 50 sec, 72°C for 1 min; followed by 35 cycles of 94°C for 30 sec, 52°C for 30 sec, 72°C for 1 min; and final extension at 72°C for 6 min and held at 10°C.

### 2.3 Mitochondrial COI sequencing and analysis

PCR products were sequenced using both forward and reverse primers, on an ABI 3730 automated sequencer at the University of Dundee Sequencing Service. Sequences were edited in Sequencher 5.4.6 (Gene Codes Corporation, Ann Arbor, MI, USA), aligned using the Muscle algorithm in Aliview v. 1.28 (Larsson, 2014) and grouped into unique haplotypes with DnaSP v. 6 (Rozas et al., 2017). Species identification was carried out using the BOLD SYSTEMS (https://boldsystems.org/index.php) and the Basic Local Alignment Search Tool (BLAST) tool provided by the National Center for Biotechnology Information (https://blast.ncbi.nlm.nih.gov/Blast.cgi).

Haplotype frequencies were calculated and used to generate minimum spanning haplotype networks in Popart version 1.7 (Bandelt et al., 1999), both to visualise comparisons between farmed and wild sampling sites in this study and to set the Nigerian samples into a global context by comparing the geographic distributions of haplotypes with 100% match with sequences in BOLD. Arlequin v. 3.5.2.2 (Excoffier and Lischer, 2010) was used to calculate summary statistics per sampling site (number of haplotypes, *Na*; number of segregating sites, *S*; haplotype diversity, *Hd*; pairwise nucleotide diversity, *pi*), along with pairwise patterns of maternal genetic differentiation (*Fst*) and hierarchical Analysis of Molecular Variance (AMOVA). To determine whether the albinos, although found in the wild, were more related to the farmed or wild sampling sites, results were compared across three AMOVAs: 1) with the albinos considered as wild samples; 2) without inclusion of albino samples; and 3) with albinos considered as farmed samples. The AMOVAS compared variation among groups (farmed vs wild), among populations within groups (among different sampling sites within either farmed or wild), and within populations (among individuals within each sampling site). We also used independent t tests to compare the mean difference between farmed and wild *C. gariepinus* when albinos were considered as wild samples, excluded from the analysis, and as farmed samples.

### 2.4 3RAD library preparation

Prior to library preparation, the DNA of each sample was standardised to a concentration of 20 ng/μl. The 3RAD library was prepared using 96 samples, including a negative control, using the published 3RAD protocol (Bayona-Vásquez et al., 2019). The DNA was digested for 1 hr at 37 °C in a reaction mix that consisted of: 1.5 µL 10x CutSmart Buffer (New England Biolabs, Inc., UK), 5.0 µL ddH_2_O, 0.5 µL of MspI at 20 U/µL (New England Biolabs, Inc., UK), 0.5 µL of BamHI-HF at 20 U/µL (New England Biolabs, Inc., UK), 0.5 µL of ClaI at 20 U/µL (New England Biolabs, Inc., UK), 1 µL 5 µM double-stranded iTru read 1 adapter, 1 µL 5 µM double-stranded iTru read 2 adapter (BadDNA, The University of Georgia, USA), and 5 µL DNA. After incubation at 37°C for 1 hour, 2.0 µL dH_2_O, 1.5 µL ATP (10 µM), 0.5 µL 10x Ligase Buffer, and 1.0 µL T4 DNA Ligase (100 units/µL, NEB M0202L buffer diluted 1:3 in NEB B8001S enzyme dilution buffer) were added to each reaction, before running the digested/adapter-ligated mixtures in a thermocycler with the following conditions: 22°C for 20 min, 37°C for 10 min, 22°C for 20 min, 37°C for 10 min, 80°C for 20 min, then hold at 10°C. The ligated products were pooled by transferring 10 µL from each row in a plate (12 wells) into a new 1.5 mL centrifuge tubes (10 µL x 12 well = 120 µL/row). From each of the 8 tubes containing 120 µL of ligated product, 60 µL was transferred into a single 1.5 mL tube to yield 480 µL (60 µL x 8) of ligation product. The remaining 60 µL from each strip were kept in the freezer for potential future use. The 480 µL pool was split into two (240 µL each) 1.5 mL tubes before purification.

Magnetic bead clean-up was performed for each pool separately using NEBNext Ultra II DNA Library Prep with Sample Purification Beads (New England Biolabs Inc., UK). The clean-up was performed at a dilution of 0.9x, followed by resuspension in 30 µL of dH_2_O. The two cleaned products were pooled into one tube (total of 60 µL). To generate full-length library constructs, cleaned ligated products were PCR amplified in a 50 µL reaction volume containing 10.0 µL Kapa HiFi Buffer (Roche, Basel, Switzerland), 1.5 µL dNTPs (10 mM), 7.5 µL ddH_2_O, 1.0 µL Kapa HiFi DNA Polymerase (1 unit/µL), 5.0 µL iTru5 primer (5 µM), and 5.0 µL iTru7 primer (5 µM), 20 µL ligated DNA fragments from the previous step (Bayona-Vásquez et al., 2019). Since the protocol utilizes 20 µL of ligated product for the PCR, we performed three reactions from the 60 µL ligation pool. In a thermocycler, the PCR master mix was amplified using the set-up: 98°C for 1 min.; 12 cycles of 98°C for 20 sec, 60°C for 15 sec, 72°C for 30 sec; 72°C for 5 min; hold at 15°C. To verify the success of the library preparation, we ran 5 µL of the PCR product with 2 µL loading dye on a 2.0% agarose gel for 45 minutes at 100 volts along with a 1kb DNA Marker (Promega, Madison, Wisconsin, USA). A successful library preparation is indicated by a bright evenly distributed DNA smear around ∼300-800 bp. All three PCR products were pooled purified with NEBNext Ultra II DNA Library Prep Sample Purification Beads in a 1:1.5 (DNA:Beads) ratio and cleaned DNA was eluted in 44 µL of ddH_2_0. The purified product was quantified using a Qubit Fluorimeter broad-range assay (Thermo Fisher Scientific, Waltham, Massachusetts, USA).

#### 2.4.1 Size selection and sequencing

The pooled library was size-selected using a Pippin Prep (Sage Science, Beverly, MA, USA) with a 2% dye-free Marker L agarose gel cassette (CDF2010), set to capture fragments of 300-450bp, and eluted in 40 µL of Tris-TAPS (N-[tris (hydroxymethyl) methyl]-3-amino propane sulfonic acid) buffer. The eluted library was quantified with the Qubit high-sensitivity assay (Thermo Fisher Scientific, Waltham, Massachusetts, USA) and sequenced (150bp paired-end) on a single lane Novaseq X (Illumina, San Diego, California, USA) at Novogene Co., Ltd (Cambridge, UK).

#### 2.4.2 3RAD sequencing analysis

Raw reads were demultiplexed into individual samples using the process_radtags module in Stacks V2.65 (Rivera-Colón and Catchen, 2022). Filtering parameters were set to drop reads with a Phred quality score of 20 or less and to remove any reads with uncalled bases. The quality of the demultiplexed reads including base quality scores, per-based sequence content, per-sequence quality scores, and sequence length distribution were assessed using FastQC (Andrews, 2010). Reads were mapped to the *C. gariepinus* reference genome GCF_024256425.1 using BWA-MEM (Li and Durbin, 2009). Population genetics analysis was performed in Stacks V2.65 (Rivera-Colón and Catchen, 2022), with parameters set to keep loci present in all the nine sampling sites (p = 9) and in at least 80% of individuals per sampling site (r = 0.80). Minimum minor allele count required to process a SNP was set at three (min-mac = 3), with SNP calls restricted to only the first per locus (-- *write-single-snp*). Summary statistics (*Ho*, *He*, *pi*) and inbreeding coefficients based on the difference between *Ho* and *He* (*Fis*) were computed for each sampling site and compared between farmed and wild sites.

#### 2.4.3 Genetic differentiation, population structure and admixture

Pairwise *Fst* was calculated between each pair of sampling sites using Genepop output from the Stacks *populations* analysis (Rivera-Colón and Catchen, 2022). To assess genetic differentiation among sampling sites, *Fst* values were calculated using the *hierfstat* (Goudet, 2005) and adegenet (Jombart et al., 2023) packages in R (R Core Team, 2018). The Genepop file was read into R and converted to a genind object to compute pairwise *Fst* values using the genet.dist function with the “WC84” method for implementation of Weir and Cockerham (1984) *Fst*. Clustered heatmaps based on the *Fst* values were generated using the Euclidean distance function from the *pheatmap* package in R (Kolde, 2019). Using the *boot.ppfst* function, a permutation test with 1,000 bootstrap replicates was performed to assess the significance of the reported *Fst* values (Goudet, 2005).

To visualise genetic variation with and between individuals in the farmed and wild groups, Principal Components Analysis (PCA) was conducted based on allele frequencies from data contained in the *Genepop* file generated from Stacks populations analysis. The *Genepop* file was loaded into R using the *Adegenet* package and converted into *GenInd* object (Jombart, 2008). The *GenInd* object was used to generate a population sample table with individual samples and population ID and converted into a dataframe. To visualise genetic variation and relatedness among individuals within, and differentiation across, sampling sites through patterns of clustering, we used the scale function *scaleGen* and *dudi.pca*. PCA eigenvalues were added to the dataframe and a scatterplot function was performed using *ggplot2* package in R (Wickham, 2016) to visualise principal components 1 and 2, which explained most of the genetic variation in the data. The *scale_color_manual* function in *ggplot2* package was used to assign unique colours to differentiate the different sampling sites in the cluster.

Admixture analysis was conducted using the ADMIXTURE software (Alexander et al., 2009). To infer the optimal number of ancestral populations (K) contributing to the genetic structure of the *C. gariepinus* sampling sites, cross-validation was performed for K values ranging from 2 to 10, using 1000 bootstrap replicates and 3 iterations. The K value associated with the lowest cross-validation error was selected as the best estimate of population structure. K = 2 was also assessed to test the hypothesis that farmed individuals can be differentiated from wild ones.

## 3 Results

### 3.1 mtDNA haplotype distribution and diversity in farmed and wild *C. gariepinus* populations

Overall, 104 COI sequences of approximately 500 to 680 bp were obtained (GenBank access numbers: PQ7861-PQ7964) and collapsed to eleven unique haplotypes; identification on the BOLD Systems and BLAST search engines returned a match for *C. gariepinus* for all, with similarity scores ranging from 99.68 to 100% (Table 2Table). The aligned sequences had a mean nucleotide composition of C=26.19%, T=28.67%, A=27.45%, and G=17.69%. All eleven haplotypes were found in fish sampled from the wild sampling sites, whereas only four haplotypes were found in samples from farms. Two farmed haplotypes (10 and 11) were shared only with the albino “wild” samples (w_DKAL). In contrast, the remaining two were shared with one wild individual from w_KDC (haplotype 5) and wild individuals from w_BYC, w_KDC and w_LGC (haplotype 7), although the latter was found in only a single farmed individual from site f_LAC. Haplotype 10 was found in the majority of the farmed samples (31 out of 43), whereas among the wild sampling sites, haplotype 8 was most common (n = 27/58), with the remaining samples distributed across different haplotypes. Haplotypes 1 – 4, 6, 8 and 9 were unique to the wild sampling sites.

**Table 2.** Haplotype identification table based on the Barcode of Life Database (BOLD) showing haplotype ID, sample sizes (N), species similarity percentage, and source classification (farmed/wild).

| **Haplotype ID** | **N** | **Farmed** | **Wild** | **Similarity** | **Source** |
| --- | --- | --- | --- | --- | --- |
| 1 | 1 | - | w_KDC (1) | 99.83 | wild |
| 2 | 4 | - | w_DKC (4) | 99.68 | wild |
| 3 | 5 | - | w_BYC (1), w_LGC (4) | 99.84 | wild |
| 4 | 1 | - | w_KDC (1) | 99.82 | wild |
| 5 | 15 | f_CMC (4), f_LAC (9) | w_KDC (1) | 100 | farmed/wild |
| 6 | 1 | - | w_LGC (1) | 99.84 | wild |
| 7 | 8 | f_LAC (1) | w_BYC (2), w_KDC (3), w_LGC (2) | 100 | farmed/wild |
| 8 | 27 | - | w_BYC (6), w_DKC (5), w_KDC (8),<br>w_LGC (8) | 100 | wild |
| 9 | 6 | - | w_DKC (5), w_KDC (1) | 100 | wild |
| 10 | 34 | f_CMC (6), f_LAC (1),<br>f_ODC (14), f_SAC (10) | w_DKAL (3) | 100 | farmed/wild |
| 11 | 4 | f_CMC (1), f_SAC (1) | w_DKAL (2) | 100 | farmed/wild |

The haplotype network comparing farmed and wild *C. gariepinus* sampling sites revealed patterns of shared maternal ancestral relationships (Figure 2). Haplotype 8 appeared to be an ancestral haplotype shared among the wild sites, with numerous connections to other wild haplotypes and the most frequently identified farmed haplotype (H10), separated by 11 mutations. Within the wild haplotypes, mutations separating the putative ancestral haplotype from the connecting haplotypes ranged from one for haplotypes 3, 5, and 9 to two for haplotype 7. The most differentiated wild haplotype (H4) was separated from H8 by 5 mutations and H7 by 3 mutations but was only found in a single individual (from w-KDC; Table 2). Farmed haplotype 11 (H11) was separated from H10 by six mutations.

**Figure 2.**
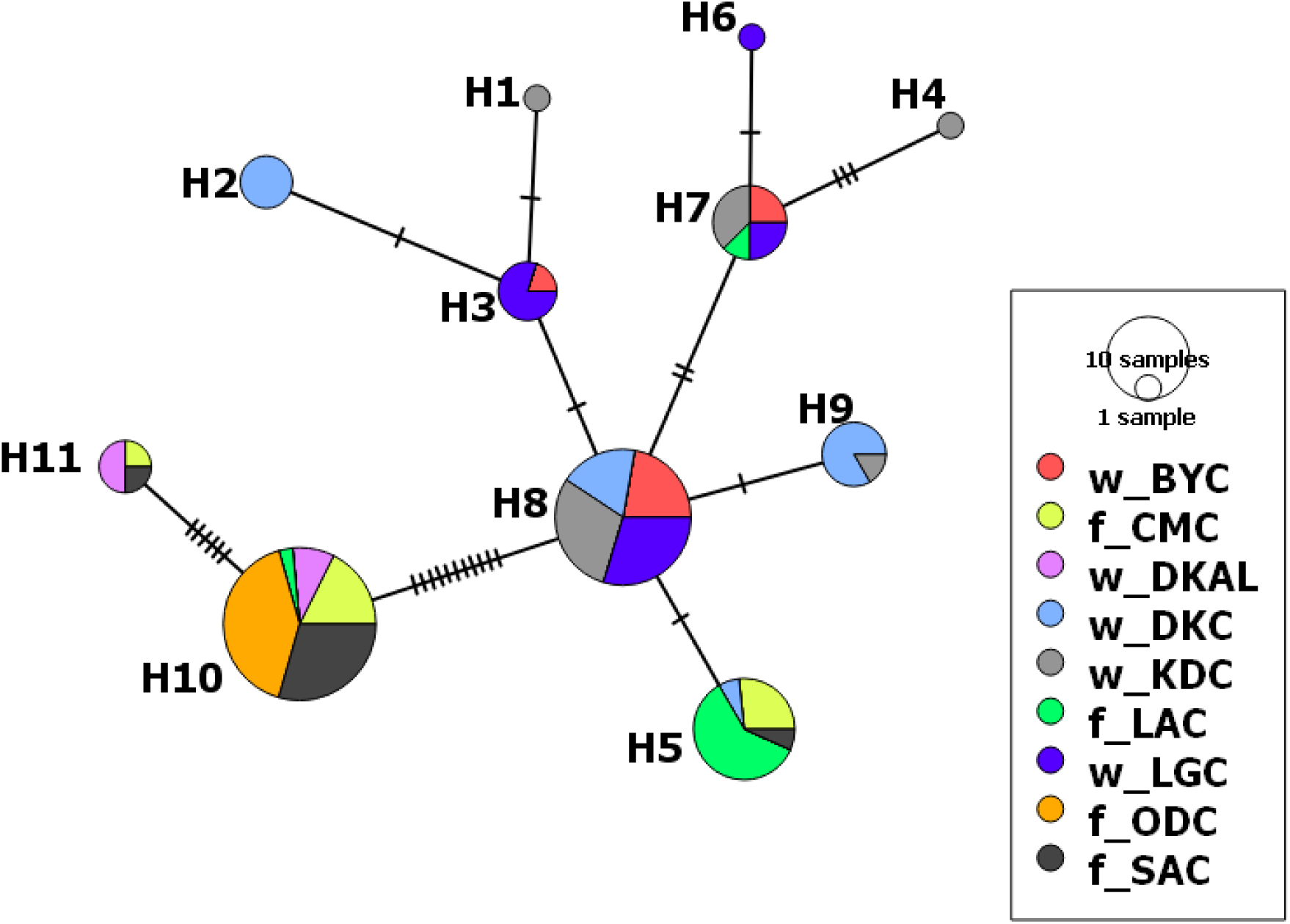
Haplotype network showing genetic relationships of farmed (f_) and wild (w_) *Clarias gariepinus* populations from Northeast (w_BYC, w_DKAL, w_DKC, w_KDC, w_LGC, and f_SAC) and Southwest (f_CMC, f_LAC, f_ODC) Nigeria. Haplotype 8, shared among w_KDC, w_LGC, w_BYC, and w_DKC, appears to be the ancestral haplotype, with the most connections to other haplotypes. The albino *C. gariepinus* from Dadin Kowa dam (w_DKAL; purple) is the only wild population which had the farmed-dominated haplotypes 10 and 11.

The BOLD Systems search results for haplotypes that had a 100% match (5, 7 – 11) revealed that these had global distributions in samples from fish farms across Africa, Asia, and the Middle East (Supplementary Table 1). Haplotypes 5, 7, and 8 were found in Nigeria, based on previous studies conducted in the country (Iyiola et al., 2018; Nwani et al., 2011). Haplotype 8, frequently found in our wild fish samples, was also found in farm samples from Israel, Thailand, Bangladesh, Syria, and India. In contrast, the frequent farmed haplotype 10 was found in DR Congo and Brazil (Supplementary Table 1).

### 3.2 mtDNA population genetic diversity

Based on the COI Sanger sequences, fish from wild sampling sites (including the albino w_DKAL) had more haplotypes (ranging from 2 – 5; average = 3.6) than those from farmed sites (ranging from 1 – 3; average = 2.5), with the highest number in site w_KDC (n = 5) and only a single haplotype in samples from f_ODC (Table 3). There was no significant difference in the number of haplotypes (Na) between the farmed and the wild groups if the albino w_DKAL samples were considered to be a wild population, nor if those samples were excluded (Table 4). However, when the albino population was included with the farmed sampling sites, there was a significance difference between farmed and wild sites (Table 4). The farmed sampling sites exhibited a non-significant trend for more segregating sites than wild sampling sites regardless of whether albinos were considered as wild, not included in the analysis, or considered as farmed (Table 4). Haplotype diversity (*Hd*) tended to be higher in wild populations (average = 0.646; highest *Hd* observed in w_DKC) compared to farmed (average = 0.321) sampling sites, except for f_CMC, which had higher *Hd* than some of the farmed sampling sites (Table 4). However, the differences were not significant, regardless of whether albino samples were considered as wild, excluded from the analysis or considered as farmed. Pairwise nucleotide diversity (*pi*) was highest in the f_CMC population and overall tended to be higher in the farmed (average = 0.005) than the wild (average = 0.003) populations, except for the albino samples, which showed *pi* more similar to the farmed sampling sites (w_DKAL = 0.006). However, differences between farmed and wild were not significant, even when the albinos were considered as farmed.

**Table 3.** Origin of farmed and wild African catfish (*Clarias gariepinus*) collected from various states in Nigeria, indicating the sampling site (Population) and code, region, the type of population (Source), number of haplotypes (Na), number of segregating sites (S), haplotype diversity (Hd) ± standard deviation (s.d), and pairwise nucleotide diversity (pi ± s.d).

| Population | Code | Region | Source | Na | S | Hd ± s.d | $\pi \pm$ s.d |
| --- | --- | --- | --- | --- | --- | --- | --- |
| CMC farm | f_CMC | southwest | Farmed | 3 | 18 | 0.618±0.104 | 0.012±0.007 |
| LAC farm | f_LAC | southwest | Farmed | 3 | 14 | 0.346±0.172 | 0.004±0.003 |
| Dutch *Clarias* | f_ODC | southwest | Farmed | 1 | - | - | - |
| SAC farm | f_SAC | northeast | Farmed | 3 | 18 | 0.318±0.164 | 0.005±0.003 |
| Kiri dam | w_KDC | northeast | Wild | 5 | 7 | 0.659±0.123 | 0.003±0.002 |
| Lake Geriyo | w_LGC | northeast | Wild | 4 | 4 | 0.667±0.099 | 0.002±0.002 |
| River Benue Yola | w_BYC | northeast | Wild | 3 | 3 | 0.556±0.163 | 0.002±0.001 |
| Dadin Kowa dam | w_DKC | northeast | Wild | 4 | 4 | 0.752±0.056 | 0.002±0.002 |
| Dadin Kowa dam | w_DKAL | northeast | Wild | 2 | 6 | 0.600±0.175 | 0.006±0.004 |

**Table 4.** Multiple t-tests comparing farmed and wild *Clarias gariepinus*. For each tested parameter results are presented for three alternative scenarios: when the w_DKAL albino population considered as wild, excluded from the analyses, or treated as farmed. The columns show statistical information including t-statistics and associated p-values, degrees of freedom (d.f.), and the 95% lower (CI lower) and upper bound (CI upper) confidence intervals.

| **Parameter** | **Farmed** | **Wild** | **t-statistic** | **p-value** | **d.f.** | **95% CI lower** | **95% CI upper** |
| --- | --- | --- | --- | --- | --- | --- | --- |
| Na (albino as wild) | 2.5 | 3.6 | -1.540 | 0.168 | 6.893 | -2.794 | 0.594 |
| Na (without albino) | 2.5 | 4.0 | -2.324 | 0.061 | 5.769 | -3.095 | 0.094 |
| Na (albino as farmed) | 2.4 | 4.0 | -2.799 | 0.027 | 6.815 | -2.959 | -0.241 |
| S (albino as wild) | 12.5 | 4.8 | 1.776 | 0.169 | 3.178 | -5.668 | 21.068 |
| S (without albino) | 12.5 | 4.5 | 1.835 | 0.157 | 3.246 | -5.295 | 21.295 |
| S (albino as farmed) | 11.2 | 4.5 | 1.831 | 0.134 | 4.468 | -3.054 | 16.454 |
| Hd (albino as wild) | 0.321 | 0.647 | -2.496 | 0.078 | 3.416 | -0.715 | 0.063 |
| Hd (without albino) | 0.321 | 0.659 | -2.548 | 0.070 | 3.598 | -0.723 | 0.047 |
| Hd (albino as farmed) | 0.376 | 0.659 | -2.357 | 0.065 | 4.971 | -0.590 | 0.026 |
| pi (albino as wild) | 0.005 | 0.003 | 0.861 | 0.443 | 3.581 | -0.005 | 0.010 |
| pi (without albino) | 0.005 | 0.002 | 1.196 | 0.316 | 3.060 | -0.005 | 0.011 |
| pi (albino as farmed) | 0.005 | 0.002 | 1.611 | 0.180 | 4.133 | -0.002 | 0.009 |

### 3.3 mtDNA COI genetic differentiation among populations

In the AMOVA analyses, regardless of whether albinos were considered as wild samples (Supplementary Table 2), excluded (Supplementary Table 3) or considered as farmed (Table 5), the comparison of groups (farmed vs wild) did not explain a significant proportion of the variation, with the majority of the variation being found among individuals within populations, followed by among populations within the farmed and wild sampling sites. The AMOVA presented in Table 6, revealed 40.83% of genetic variability among the population associated with high genetic differentiation (*Fst* = 0.408). However, most of the genetic variation was found within the population (59.17%).

**Table 5.** Analysis of Molecular Variance (AMOVA) of mtDNA COI sequences in nine farmed and wild *C. gariepinus* populations from northeastern and southwestern Nigeria including albino as farmed samples

| Source of Variation | d.f. | SS | Var | % Var |
| --- | --- | --- | --- | --- |
| Among groups | 2 | 10.338 | -0.036 Va | -8.59 |
| Among populations within groups | 8 | 18.083 | 0.170 Vb | 41.12 |
| Within populations | 200 | 55.702 | 0.279 Vc | 67.47 |
| Total | 210 | 84.123 | 0.413 | 100 |

Notes: SS, sum of squares; Var, variance component; Va, variance components among populations; Vb, variance components among populations within groups; Vc, variance components within populations

**Table 6.** Analysis of molecular variance (AMOVA) of mtDNA COI sequences in farmed and wild *C. gariepinus* populations from the study areas.

| Source of Variation | d.f. | SS | Var | % Var | $F_{st}$ |
| --- | --- | --- | --- | --- | --- |
| Among populations | 8 | 18.083 | 0.172 Va | 40.83 | 0.408 |
| Within populations | 97 | 24.209 | 0.250 Vb | 59.17 |  |
| Total | 105 | 42.292 | 0.422 | 100 |  |

Notes: SS, sum of squares; Var, variance component; Va, variance components among populations; Vb, variance components among populations within groups; Vc, variance components within populations

### 3.4 3RAD reads summary

The demultiplexed library resulted in 1,583,447,229 reads, ranging from 1,198,636 to 35,931,402 per individual. All samples passed the FASTQ check, with a sequence per base quality score ≥ 28. The mean read coverage was 109.5x (Supplementary Figure 1a) but with varying distribution of reads across sampling sites (Supplementary Figure 1b). A total of 1,663,643 loci were assembled, ranging from 37,000 to 100,000 loci per sample (Supplementary Figure 2a). The mean rate of missing variant sites (i.e. the average percentage of genomic locations that do not have any sequence information) within sampling sites was less than 0.05% for most samples, except for one sample in the w_DKC sampling site with about 25% missing variant sites (Supplementary Figure 2b). This sample was dropped from subsequent analysis.

### 3.5 Population summary statistics

Based on 14,410 variant sites genotyped from 20,126 loci, wild samples tended to show higher maximum values and broader ranges for *Ho* (0.109 – 0.165), *He* (0.111 – 0.216), and *pi* (0.125 – 0.225) compared to the farmed sites (*Ho* = 0.118 – 0.147, *He* = 0.112 – 0.144, *pi* = 0.117 – 0.151) (Table 7). However, there was no significant difference in average *Ho* between farmed (*Ho* = 0.135) and wild (*Ho* = 0.138) sites (*t* = -0.258, *df* = 6.753, *p* = 0.804). Likewise, average *He* and *pi* did not significantly differ between farmed (*He* = 0.127; *pi* = 0.133) and wild (*He* = 0.172; *pi* = 0.183) sites (*t* = -2.277, *df* = 5.012, *p* = 0.072; *t* = -2.511, *df* = 5.148, *p* = 0.052). These results did not change when albino samples were excluded from the analyses. However, when albino samples were considered as farmed, there were significant differences between farmed and wild samples for all summary statistics except for *Ho*, where the difference between farmed (*Ho =* 0.130) and wild (*Ho =* 0.145) was not significant (*t* = -1.502, *df* = 6.863, *p* = 0.178). Average *He* and pi were significantly lower in farmed (He = 0.124; *pi* = 0.132) compared to wild (*He* = 0.188; *pi* = 0.198) samples (*t* = -4.175, *df* = 4.152, *p* = 0.013; *t* = -4.118, *df* = 3.911, *p* = 0.015 respectively).

**Table 7.** Summary of observed (*Ho*) and expected heterozygosity (*He*), nucleotide diversity (*pi*), and inbreeding coefficients (*Fis*) estimated from 14,410 variant sites genotyped from 20,126 loci, showing the genetic diversity in farmed (f_) and wild (w_) *Clarias gariepinus* in Nigeria

| Pop ID | Code | N | $H_o$ | $H_e$ | $\pi$ | $F_{is}$ |
| --- | --- | --- | --- | --- | --- | --- |
| CMC farm | f_CMC | 10 | 0.147 | 0.144 | 0.151 | 0.017 |
| Lagos farm | f_LAC | 12 | 0.142 | 0.121 | 0.127 | -0.031 |
| Ogun Dutch | f_ODC | 12 | 0.132 | 0.112 | 0.117 | -0.034 |
| SAC | f_SAC | 11 | 0.118 | 0.130 | 0.137 | 0.052 |
| River Benue | w_BYC | 9 | 0.144 | 0.205 | 0.220 | 0.196 |
| Dadin Kowa Alb | w_DKAL | 5 | 0.109 | 0.111 | 0.125 | 0.038 |
| Dadin Kowa | w_DKC | 11 | 0.165 | 0.216 | 0.225 | 0.163 |
| Lake Geriyo | w_LGC | 12 | 0.137 | 0.155 | 0.162 | 0.076 |
| Kiri Dam | w_KDC | 8 | 0.133 | 0.174 | 0.184 | 0.160 |

In terms of inbreeding, the farmed f_ODC and f_LAC samples had a slight excess of heterozygosity (i.e. negative inbreeding coefficient, *Fis*), whereas all the others were positive. For wild samples, there was only substantial evidence of inbreeding in the w_BYC, w_KDC and w_DKC sites (values of *Fis* >0.1); the albino (w_DKAL) samples showed the lowest inbreeding coefficient (*Fis =* 0.038) among the wild sites. Even when albino samples were considered as wild, inbreeding coefficients were significantly lower in farmed (*Fis* = 0.001) compared to wild (*Fis* = 0.127) sites (*t* = - 3.472, *df* = 6.707, *p* = 0.011). When albino samples were considered as farmed, the average increased for both farmed (*Fis* = 0.008) and wild (*Fis* = 0.149) sites but the difference remained significant (*t* = -4. 519, *df* = 5.577, *p* = 0.005).

### 3.6 Genetic differentiation and admixture between farmed and wild populations

The pairwise *Fst* comparisons between sampling sites from the 3RAD genotypic data showed lower genetic differentiation among the wild sampling sites (0 – 0.26) compared to the farmed sites (0.16 - 0.28) (**Error! Reference source not found.** 3). Pairwise *Fst* comparisons among wild sampling sites were significant except the comparison between w_BYC and w_KDC (*Fst* [95% bootstrapped CI] = -0.00 [- 0.003–0.002]). *Fst* was also significant for paired comparisons among the farmed sampling sites. Comparison between the albino sample (w_DKAL) and the farmed sampling sites showed consistently low but significant *Fst* values, ranging from 0.16 (95% bootstrapped CI = 0.150–0.168) when w_DKAL was compared to f_CMC to 0.28 (CI = 0.264–0.286) when compared to f_LAC. However, *Fst* values were consistently higher when the albino sample was compared to other wild sites, ranging from 0.33 (CI = 0.383–0.400) when comparing w_DKAL to w_DKC to 0.42 (CI = 0.407–0.430) when compared to w_LGC. The lowest *Fst* (-0.00) among the wild sampling sites was observed between Kiri Dam (w_KDC) and River Benue Yola (w_BYC). Hierarchical clustering analysis based on the Fst heatmap (Figure 3) indicated one large cluster consisting of the farmed sites together with the albino w_DKAL site and a second cluster containing the remainder of the wild sampling sites. The figure also indicates low genetic differentiation among wild sampling sites in Adamawa state (w_BYC, w_KDC, and w_LGC) ranging from *Fst =* -0.00 (between w_BYC and w_KDC) to *Fst =* 0.03 (between w_BYC and w_LGC). This contrasts with much higher genetic differentiation when these Adamawa sampling sites were compared with the w_DKC site in Gombe state (*Fst* values ranging from 0.10.

**Figure 3.**
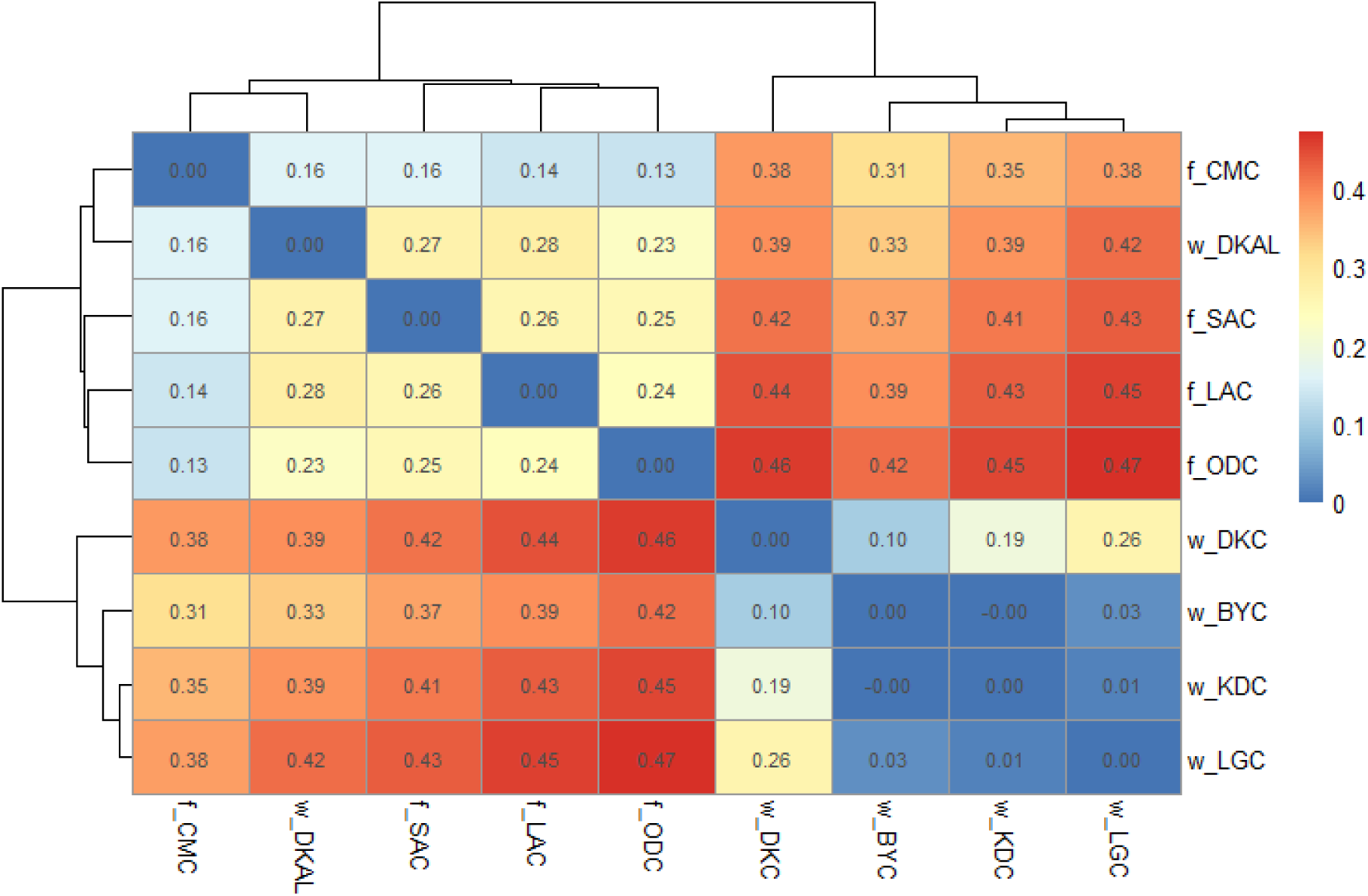
Pairwise genetic differentiation (*Fst*) among nine *C. gariepinus* from farmed and wild populations based on SNP data. f_CMC: CMC farm, f_ODC: Dutch Clarias, f_LAC: Lagos farm, f_SAC: SAC farm, w_BYC: River Benue Yola, w_LGC: Lake Geriyo, w_DKAL: Albino Dadin Kowa, w_DKC: Dandin Kowa dam, w_KDC: Kiri dam. The dendrogram displays the structure of genetic similarity among sampling sites, with more closely related sampling sites clustering together on nearby branches. The heatmap shows the degree of genetic differentiation between two sampling sites increasing from very low (blue) to very high (red) genetic differentiation.

The PCA plot also showed a clear separation between the wild and farmed samples, with principal components 1 (PC 1) and 2 (PC 2) explaining 20.7% and 13.3% of the variance, respectively (Figure 4). The distribution of individuals and sampling sites on the PCA plot showed a major cluster of all the farmed samples along the PC1 axis, but with the albino samples (green triangle) again clustering with them rather than with wild samples. Among the wild sampling sites, samples from Adamawa state (w_BYC, w_KDC, and w_LGC) clustered at the bottom (i.e. lower values of PC2) while the Gombe samples (w_DKC: Dadin Kowa dam) were distributed at the top of the PC 2 axis.

**Figure 4.**
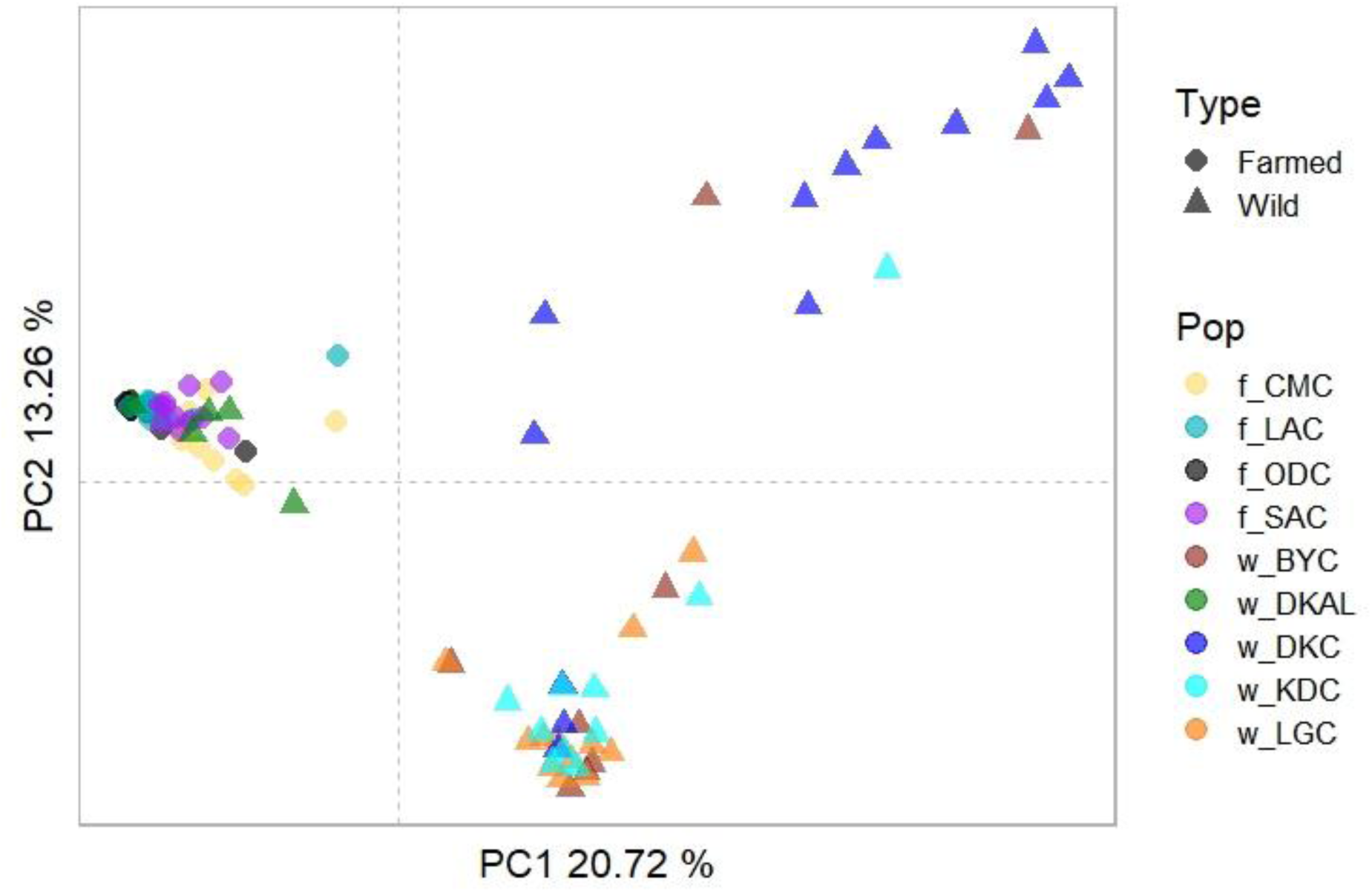
A Principal Component Analysis (PCA) plot of farmed and wild *C. gariepinus* generated from SNP data. The x-axis and y-axis represent the first and second principal components, respectively, which are linear combinations of SNP alleles that capture the maximum amount of variation in the data. PC1 (explaining 20.72% of the variation) separates the wild samples (triangles) from the farmed samples (circles), except for the wild albino population (w_DKAL) sampled from Dadin Kowa dam. PC2 (explaining 13.26% of the variance) separates some of the wild populations from one another; note that w_BYC is found in both main wild clusters.

For the admixture analysis, at K = 2 there was obvious differentiation between the farmed and wild sampling sites but again, the albino wild samples (w_DKAL) clustered with the farmed groups (Figure 5). The lowest cross-validation error value suggested that K = 5 best described the population structure. At this value, farmed f_SAC from northeast Nigeria was differentiated from their farmed counterparts from the southwest. Within the southwest populations, the CMC farm (f_CMC) was genetically distinct from the Dutch *Clarias* (f_ODC) but all individuals from f_LAC showed evidence of admixture including all of the farmed clusters. Some of the individuals from f_SAC also showed admixture with the other farmed clusters. The albino wild samples (w_DKAL) clearly clustered with Dutch *Clarias* (f_ODC), with little evidence of admixture at either of these sites. For the wild samples, as with the PCA results, the Adamawa sampling sites (w_BYC: River Benue Yola, w_LGC: Lake Geriyo, and w_KDC: Kiri dam) appeared to be differentiated from the Gombe population (w_DKC: Dadin Kowa dam). However, there was evidence of admixture within and between w_BYC and w_DKC, with some individuals from w_KDC also showing admixture with the main cluster in w_BYC. W_DKC also showed some evidence of admixture with farmed clusters.

**Figure 5.**
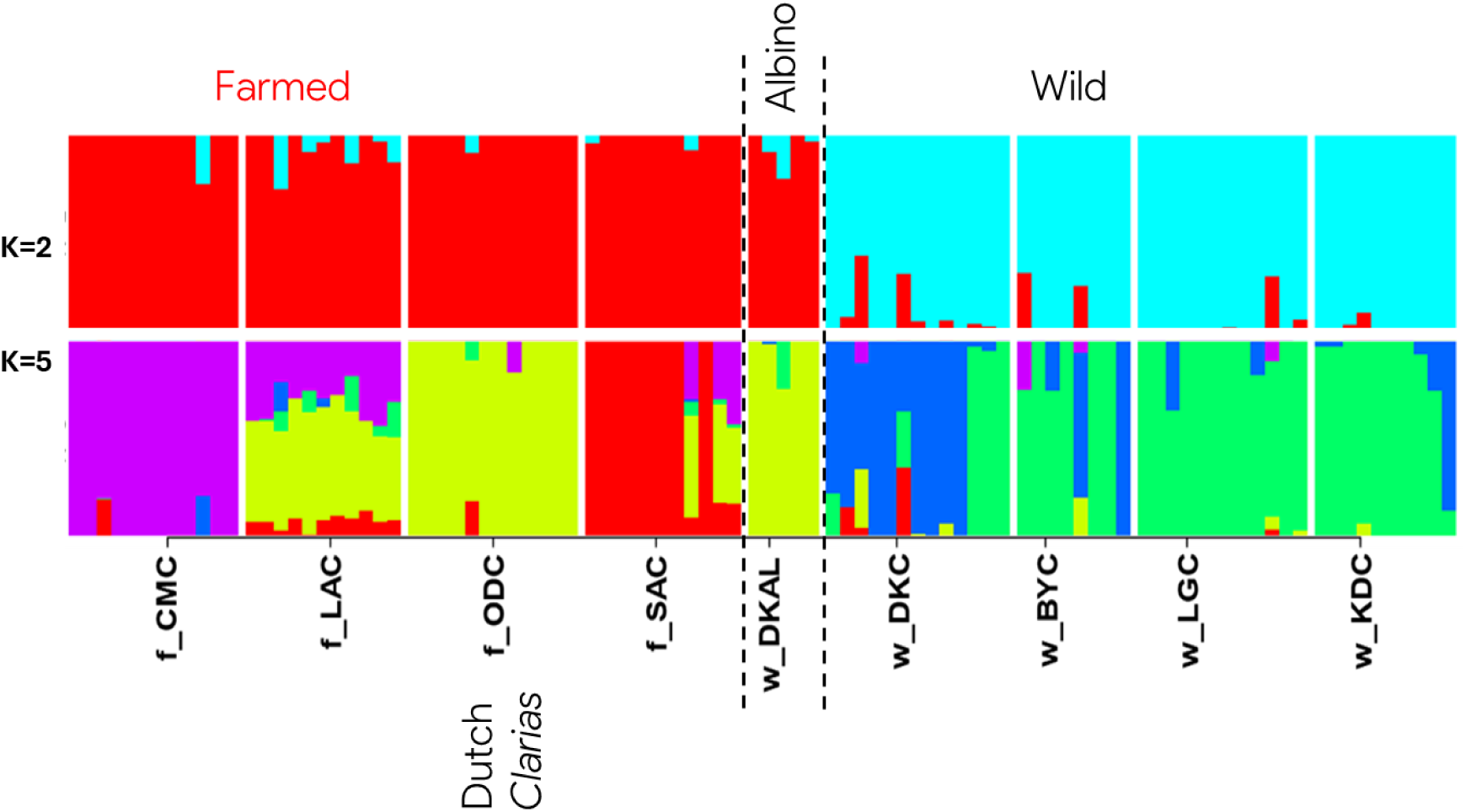
Admixture plot illustrating the genetic ancestry of individual fish samples from farmed and wild *C. gariepinus* populations based on SNP data. Each vertical bar in the plot represents an individual fish sample, partitioned into segments corresponding to inferred genetic clusters. The plot is grouped according to farmed (the first four), albino and wild (the last four). At K=2, we tested the hypothesis whether samples can be differentiated into farmed and wild sampling sites. K = 5 shows further genetic differentiation among sampling sites and hybridisation across the farmed and wild sampling sites.

## 4 Discussion

In this study, the genetic diversity and differentiation of farmed and wild sampling sites of *C. gariepinus* in Nigeria were investigated using the mtDNA COI gene and 3RAD sequencing approaches. Overall, we found that, except for an aberrant population of albino individuals captured in the wild but almost certainly of farmed origin, there were clear patterns of differentiation, and little evidence of admixture, between farmed and wild sampling sites. This suggests limited interbreeding, despite sharing of some mtDNA haplotypes, which could explain why the hierarchical AMOVA did not indicate significant variance explained by the farmed vs wild grouping. Studies have reported high genetic morphological variations in farmed and wild *C. gariepinus* populations from different water bodies. Ola-Oladimeji et al. (2017) in a study involving four wild *C. gariepinus* populations in Nigeria using morphometric and meristic methods, observed distinctive morphometric traits among the four populations. Popoola et al. (2014) while investigating genetic variation in farmed and wild *C. gariepinus* population in Nigeria, found that extensive amount genetic variation in both farmed and wild population with 99% of the total genetic variation (99%) residing within populations. There also appeared to be encouraging levels of genetic variation within the wild sampling sites, although these had greater evidence of inbreeding than in the farmed sites. Similarly, in a genetic diversity and population structure of farmed and wild *C. gariepinus* in Kenya, Barasa et al. (2017) found more numbers of haplotypes and haplotype diversities in the wild compared to farmed sites as well as a high genetic differentiation indicating genetically distinct populations. This is consistent with the admixture analyses, which suggested that farmed sites often included fish from multiple sources. Since pairwise genetic differentiation (*Fst)* based on the 3RAD data was lower among the wild than the farmed sites, it also could be a sign of bottlenecks in the wild populations. However, there was still some evidence of admixture among wild sites, which could suggest that there has not yet been substantial isolation of populations that could pose severe risks to population viability. Nevertheless, our findings that albino catfish sampled from the wild not only shared two of the farmed mtDNA haplotypes but also clustered with the farmed sampling sites in both PCA and admixture analyses of the 3RAD data, even though other normally-pigmented individuals from the same wild sampling site were highly distinctive, suggests that there are ongoing risks to the genetic integrity of wild populations unless there are measures implemented to regulate escapes. The shared genetic similarities observed between the albino and the Dutch *Clarias*, along with the presence of haplotypes found in the albino samples in DR Congo (Sonet et al., 2019), which is one of the donor countries of *C. gariepinus* to Belgium and the Netherlands where Dutch *Clarias* was developed (Holčík, 1991; Huisman and Richter, 1987), could confirm the identity of the albino samples as escaped farmed Dutch *Clarias.* We thus recommend implementing preventive measures to minimise fish escapes and enacting policies that will mandate conducting risk assessment before introducing non-native species.

### 4.1 Distribution of mitochondrial haplotypes in farmed vs wild catfish

The mtDNA analyses revealed haplotypes that were only found in wild sampling sites, suggesting limited gene flow between farmed and wild sampling sites. Only two of the haplotypes (8 and 9) unique to wild populations had a 100% match in the BOLD database, but representative samples were distributed across the Middle East and Asia; for example, haplotypes (8 and 9) have previously been found in a survey of freshwater fish diversity in Israel (Tadmor-Levi et al., 2023). This suggests a broader geographic distribution of these haplotypes in the Middle East and Asia. In contrast, all of the haplotypes obtained from our sampled fish farms had also been described from other regions, which could reflect shared sources of farmed fish. Haplotype 5, most commonly found in farmed sampling sites (f_LAC and f_CMC) but also one wild site (w_KDC), was previously reported in the wild in southeastern Algeria (Behmene et al., 2022) and southeast Nigeria (Nwani et al., 2011). The presence of Haplotype 5 in predominantly farmed sampling sites suggests a possible collection of local wild fish. This is a common practice among catfish farmers in Nigeria, who often acquire gravid broodstock from fishers during the peak rainy season when wild *Clarias* spp. are about to begin breeding (MKS, personal observation). Purchasing broodstock from the wild might be preferred due to its cost-effectiveness compared to buying from farms, if breeders view wild-caught broodstock as more economically viable. This decision to buy wild-caught broodstock is therefore not driven by genetic considerations (Ibiwoye, 2018). Similarly, predominantly wild haplotype 7, which was also found in farmed f_LAC, has in previous studies been reported to be a wild haplotype in both North central (Iyiola et al., 2018) and Southeastern Nigeria (Nwani et al., 2011). The presence of predominantly farmed haplotypes 10 and 11 in the albino samples (w_DKAL) reinforces questions about the true wild status of the w_DKAL samples. The absence of haplotypes 10 and 11 in previous DNA barcoding studies conducted in Nigeria, which primarily focused on wild samples, suggests that these haplotypes were not previously documented in sequences submitted to BOLD from Nigerian studies. However, both haplotypes 10 and 11 were identified in wild catfish from the Lower and Middle Congo Rivers, as well as in three major drainage basins of the Lower Guinean ichthyological province, namely Kouilou-Niari, Nyanga, and Ogowe (Sonet et al., 2019).

Pairwise nucleotide diversity of mtDNA was higher in some of the farmed than the wild populations, possibly because the farmed samples had come from a variety of sources and so included differentiated haplotypes (indicated by the higher number of segregating sites). This was also suggested by the high admixture in the 3RAD analyses for some of the farmed populations. Previous studies have confirmed that the farmed haplotypes are sourced from the wild as reported in Nigeria (Iyiola et al., 2018; Nwani et al., 2011), Algeria (Behmene et al., 2022), and DR Congo (Sonet et al., 2019).

### 4.2 mtDNA genetic diversity and differentiation

The mtDNA COI result suggests that wild sampling sites overall harbour a greater diversity of mtDNA haplotypes, which may reflect their larger effective population sizes than on fish farms, which typically use relatively few individual fish as broodstock. Nevertheless, the overall genetic diversity parameters (*Hd* and *pi*) obtained in this study are very low compared with results from previous studies. For example, Kundu et al. (2023) using mtDNA COI to study wild *C. gariepinus* from the Nyong River in Cameroon reported 20 haplotypes with 81 segregating sites, haplotype diversity (*Hd*) = 0.854, and nucleotide diversity (*pi*) = 0.258. A similar study conducted in three rivers in southwest Nigeria using the mtDNA *cytb* gene revealed 53 haplotypes, with greater haplotype diversity (*Hd* = 0.999) and nucleotide diversity (*pi* = 0.073) than the current study (Popoola, 2022). Nyunja et al. (2017) studied five *C. gariepinus* hatchery populations in Kenya, and found 33 haplotypes, 60 segregating sites, high haplotype diversity (*Hd* = 0.988 ± 0.031), and low nucleotide diversity (pi = 0.024 ± 0.026). The relatively high genetic diversity (*Hd* = 0.618 ± 0.104, *S* = 18) found in the CMC farm can be explained by the periodic introduction of new broodstock into the hatchery from multiple sources (Ibiwoye, 2018). Alal et al. (2021) suggested that the genetic diversity in small *C. gariepinus* populations in the wild as observed in Lake Kenyatta, Kenya, can be improved through stock augmentation by conservation scientists. For example, this approach could be used to boost declining *C. gariepinus* in lakes due to overfishing and periodic drying. A shared haplotype was observed between one individual from a farm (f_LAC) and wild fish from three sampling sites, suggesting that broodstock for the f_LAC farm may have been sourced from the wild. Interestingly, shared haplotypes between farmed and wild samples were observed in Kenya farmed *C. gariepinus* and wild samples from Lakes Victoria, Kanyaboli, and Baringo (Barasa et al., 2017).

The AMOVA analysis showed high genetic differentiation among the nine populations (global *Fst* = 0.408), but that the percentage of genetic variation was mostly explained by differences among populations within farmed and wild categories, as well as within populations themselves, rather than differences between farmed and wild populations. This result was the same, regardless of how the albino population was considered but the absence of shared haplotypes between the albino and normally pigmented wild catfish captured at the same site (Dadin Kowa dam) is noteworthy. This result was similar to the study involving farmed and wild *C. gariepinus* in Nigeria reported by Suleiman et al. (2020), who observed 4% of the variance among the populations and 96% within the population. Popoola (2022) in a study of three wild *C. gariepinus* populations in Nigeria, also found a high genetic differentiation among populations (Fst = 0.757), however, contrary to our findings, most of the genetic variation was found among populations (75.73%) against 24.27% found within populations. The closer genetic relationship of the albino fish to farmed fish from other sites rather than to the wild normally-pigmented samples caught at the same site suggests that their escape occurred relatively recently. Other studies using mtDNA markers have been conducted to investigate the genetic differentiation among *C. gariepinus* populations. For example, Alal et al. (2021) in their study of lacustrine and riverine *C. gariepinus* populations in Kenya using the mtDNA D-loop, observed distinct genetic patterns and population structures among lacustrine and riverine populations, with lower but significant genetic differentiation (*Fst* = 0.166, p < 0.001) among populations than found in the current study. In a similar study using the cytb gene, Popoola (2022) found high genetic differentiation (*Fst* = 0.75 – 0.95) among three Southwest populations in Nigeria. Collectively, these studies have shown that the COI gene and other mtDNA markers including the D-loop and cytb can be applied to successfully differentiate different populations of *C. gariepinus*. This cheap molecular approach compared to deep sequencing could be a vital tool for the conservation of *C. gariepinus* in Nigeria.

### 4.3 3RAD genetic diversity in farmed and wild populations

The genome-wide SNP data show consistently higher genetic diversity in the wild compared to farmed samples, a finding that is concordant with the mtDNA COI genetic diversity results. However, the reduced genetic diversity in farmed sites may be due to selective breeding and limited broodstock, as has been demonstrated in other farmed fish species (Zeinab et al., 2014). Maintaining genetic diversity in hatchery populations requires the farmer to have genetically diverse base broodstock populations (Fernández et al., 2014). However, one of the challenges facing the catfish industry in Nigeria is access to quality broodstock from reliable hatcheries (Adeleke et al., 2021). The control of artificial reproduction in hatcheries is an important management practise critical for successful spawning and obtaining of good quality eggs (Duncan et al., 2013). Indiscriminate breeding practices among fish farmers, however, makes broodstock management difficult, and is detrimental to maintaining genetic diversity in Nigerian fish farms (Sanda et al., 2024). .

Despite greater genetic diversity among wild sampling sites compared to farmed ones, it is still low when compared to results obtained from other studies of African catfish that have been conducted using other types of nuclear markers. For example, values for *Ho* (0.109 – 0.165) and *He* (0.111 – 0.216) in the current study were less than was found in catfish samples from Asejire dam in Nigeria (*Ho = 0.*409, *He* = 0.566) using microsatellite markers (Awodiran et al., 2019). In a similar study that used microsatellite markers to explore the genetic diversity of *C. gariepinus* hatchery populations across northeastern and central Thailand, Wachirachaikarn and Na-Nakorn (2019) noted substantial heterozygosity, with values ranging from 0.52 to 0.72 for *Ho* and 0.67 to 0.77 for *He*. Several factors including overexploitation, environment change, pollution, and habitat destruction have been identified among the main causes of population decline that are increasingly threatening fish diversity in the wild (Martinez et al., 2018; Petit-Marty et al., 2022). These factors were noticeable in some of the wild sampling sites in the present study where the species appeared to be depleted, such as the River Benue Yola, which is affected by seasonal drying during the extended dry season period when water levels in Nigerian rivers and lakes decrease substantially. The review by Emmanuel et al. (2021) highlighted how overexploitation, habitat fragmentation, and pollution are posing threats and endangering fish diversity in Nigeria. Improving fisheries management and regulation of fishing activities including habitat restorative programmes are therefore important for maintaining genetic diversity of wild *C. gariepinus*.

The observed higher *Fis* in the wild compared to farmed *C. gariepinus,* is consistent with results obtained in a study of farmed and wild *C. gariepinus* in Nigeria using RAPD markers (Suleiman et al., 2020). Meanwhile the excess heterozygosity observed in two of our farmed sites (f_LAC and f_ODC) aligns with a study of farmed and wild *C. gariepinus* in southwest Nigeria using RAPD and microsatellite markers (Awodiran and Afolabi, 2018), which reported negative *Fis* in one of the farmed sites, suggesting excess heterozygosity compared to those that had positive *Fis*. The high *Fis* in the wild sampling sites in the present study could also be due to declining population sizes leading to an increase in the likelihood of mating between related individuals. This was demonstrated in the giant fish *Arapaima gigas* in the Amazon, which experienced a loss of genetic diversity and high inbreeding rates caused by illegal fishing and environmental pressure reducing the effective population size (Fazzi-Gomes et al., 2021). Inbreeding in wild *C. gariepinus* could be reduced by promoting sustainable fishing and ensuring that conservation efforts take into account the importance of maintaining genetic diversity and promoting gene flow through habitat connectivity.

### 4.4 Genetic differentiation among farmed and wild populations

Overall, there was consistently high genetic differentiation between the farmed and wild *C. gariepinus* sampling sites, suggesting that mixing between farmed and wild samples has so far been limited. However, there is consistent genetic evidence that the albino fish (w_DKAL), while captured in the wild, had escaped from a farm. This shared genetic similarity between the albino fish and farmed sampling sites is consistent with results in similar studies where farmed escapes show higher similarity to their source populations (Erkinaro et al. (2010); (Šegvić-Bubić et al., 2016). Additionally, using Bayesian inference to analyse a 14-year dataset of farmed escapees from 54 rivers in western Norway, Mahlum et al. (2021) concluded that the abundance of farmed escapees in rivers was correlated with aquaculture intensity. If the albino sample is considered as farmed, it can be concluded based on the first admixture cluster of K = 2, and the PCA that *C. gariepinus* populations in Nigeria, despite their low genetic diversity, can be differentiated according to farmed and wild using the genome-wide SNPs. The significant *Fst* values between farmed and wild sampling sites is an important indicator for the implementation of conservation strategies that will manage these different farmed and wild sites separately to preserve their unique diversity. Relevant scientific data such as this are relevant to the management of fish and aquaculture activities (Kemp et al., 2023).

The low *Fst* among wild northeast sampling sites from Adamawa (w_BYC, w_KDC, and w_LGC) compared to substantial genetic differentiation from another northeast sampling site from Gombe state (w_DKC) provides evidence for geographical differentiation within the northeast region. However, *Fst* among the wild sampling sites was consistently lower than among the farmed sites. The PCA and admixture analyses also suggest that there has been some mixing between both farmed and wild populations, which could reflect ongoing gene flow or transport of samples between sites. The admixture in the wild sites could be linked to the connectivity that Lake Geriyo and Dadin Kowa dam share with River Benue in the northeast. Lake Geriyo is a tributary to River Benue while Dadin Kowa dam is located on the Gongola River, which is a major tributary of the Benue River (Essien et al., 2019; Hassan et al., 2015). Connectivity either from the main river itself or from tributaries, most especially in the downstream direction, facilitates movement of fish and genetic material between different sites, leading to admixture (Meulenbroek et al., 2024).

The movement of farmed fish, including broodstock and fingerlings from different sources, has also been reported as a common practice among farmers in Nigeria (Ibiwoye, 2018). While some broodstock are collected from private fish farms (Mekuleyi et al., 2022), it is not uncommon to source them from natural water bodies like lakes and rivers (Eze et al., 2022). The choice of whether to source from a farm or the wild would likely depends on factors such as availability, cost, and desired genetic trait (Eyo et al., 2021). Overall, the *Fst* values observed in this study were higher than results obtained from other *C. gariepinus* studies using different markers including microsatellites, possibly because other studies focused only on wild populations. For example, in an assessment of genetic differentiation of *C. gariepinus* from three regions in India using microsatellites, Ezilrani and Christopher (2015) observed low to moderate *Fst* (0.159 – 0.200) values that were comparable to those observed between farmed sampling sites in this study (*Fst* = 0.16 – 0.25). This suggests that even after years of *C. gariepinus* farming in Nigeria, there is still a noticeable degree of genetic differentiation maintained across different farm and wild sites.

## 5 Conclusions

A combination of traditional mtDNA Sanger sequencing and genome-wide SNP analyses based on 3RAD sequencing has illustrated the importance of applying genetic and genomic tools to the management and conservation of farmed and wild fish species. Specifically, we present evidence for the current relative genetic integrity of wild populations, but also forensic evidence of past farm escapes and mixing between wild and farmed populations. These methods also provide proxies for genetic health such as relative levels of inbreeding, which provide valuable insights for the formulation of conservation and management strategies aimed at preserving wild fish populations and regulating aquaculture activities to mitigate the risk of escape. The evidence of low genetic diversity in wild Nigerian catfish and escapes from fish farms calls for management actions including the implementation of genetic monitoring, habitat restoration, and sustainable aquaculture and fishing practices. Future research directions should consider sampling across more geographical locations and farms to further understand the dynamics of genetic diversity and population structure in farmed and wild fish.

## Supporting information

https://github.com/mksanda/Clarias_gariepinus_3RAD

## DATA AVAILABILITY STATEMENT

Raw sequence data were deposited in the European Nucleotide Archive (ENA) SRA (PRJEB78629). Mitochondrial DNA cytochrome c oxidase 1 (CO1) sequences have been deposited to GenBank, with accession numbers PQ197861-PQ197964. All additional data including scripts and metadata are available at https://github.com/mksanda/Clarias_gariepinus_3RAD.

## ACKNOWLEDGEMENTS

The authors are grateful to Bitrus Philip from the Department of Fisheries Gombe for providing the albino and other samples from Dadin Kowa, and Chris Sanda, Pius Sanda, and Lisa Sanda for helping with the Northeast sampling. Thank you to Sunday Unwana for your help during the sampling period in the Southwest region of Nigeria and for interpreting to the locals the purpose of our visit. We would like to acknowledge the generous funding from the Society of Systematic Biologists that allowed MKS to attend the RADCAMP2023 edition at Columbia University, New York to learn the 3RAD approach, and to all the organisers and instructors, including Natalia Bayona Vasques who was still providing support after the workshop. Sandra Hoffberg, Daren Eaton, Isaac Overcast, and Edgar Benavides provided great genomics training and bioinformatic analysis of 3RAD data.

## REFERENCES

Adeleke, B., Robertson-Andersson, D., Moodley, G. & Taylor, S. 2021. Aquaculture in Africa: A comparative review of Egypt, Nigeria, and Uganda vis-a-vis South Africa. Reviews in Fisheries Science & Aquaculture, 29, 167–197. 10.1080/23308249.2020.1795615

Alal, G. W., Barasa, J. E., Chemoiwa, E. J., Kaunda-Arara, B., Akoll, P. & Masembe, C. 2021. Genetic diversity and population structure of selected lacustrine and riverine populations of African catfish, Clarias gariepinus (Burchell, 1822), in Kenya. Journal of Applied Ichthyology, 37, 427-438. 10.1111/jai.14167

Alexander, D. H., Novembre, J. & Lange, K. 2009. Fast model-based estimation of ancestry in unrelated individuals. Genome Research, 19, 1655–64. 10.1101/gr.094052.109

Andrews, S. 2010. FastQC: a quality control tool for high throughput sequence data. Available online at: http://www.bioinformatics.babraham.ac.uk/projects/fastqc.

Ataguba, G. A., Annune, P. A. & Ogbe, F. G. 2009. Induced breeding and early growth of progeny from crosses between two African clariid fishes, Clarias gariepinus (Burchell) and Heterobranchus longifilis under hatchery conditions. Journal ofApplied Biosciences, 14, 755–760.

Awodiran, M., Adeniran, F., Akinwale, R. & Akinwande, A. 2019. Microsatellite variability of two populations of *Clarias gariepinus* (Siluriformes, Clariidae) in Nigeria. Vestnik Zoologii, 53, 195–208. 10.2478/vzoo-2019-0020

Awodiran, M. O. & Afolabi, O. 2018. Genetic diversity in cultured and wild population of *Clarias gariepinus* (Burchell, 1822) in Nigeria using Random Amplified Polymorphic DNA (RAPD) and microsatellite DNA. Fisheries and Aquaculture Journal, 9, 1–6. 10.4172/2150-3508.1000247

Baird, N. A., Etter, P. D., Atwood, T. S., Currey, M. C., Shiver, A. L., Lewis, Z. A., Selker, E. U., Cresko, W. A. & Johnson, E. A. 2008. Rapid SNP discovery and genetic mapping using sequenced RAD markers. PLoS One, 3, e3376. 10.1371/journal.pone.0003376

Baldwin, C. C., Mounts, J. H., Smith, D. G. & Weigt, L. A. 2009. Genetic identification and color descriptions of early life-history stages of Belizean *Phaeoptyx* and *Astrapogon* (Teleostei: Apogonidae) with Comments on identification of adult Phaeoptyx. Zootaxa, 1, 1–22. 10.11646/zootaxa.2008.1.1

Bandelt, H. J., Forster, P. & Röhl, A. 1999. Median-joining networks for inferring intraspecific phylogenies. Molecular Biology and Evolution, 16, 37–48. 10.1093/oxfordjournals.molbev.a026036

Barasa, J. E., Abila, R., Grobler, J. P., Dangasuk, O. G., Njahira, M. N. & Kaunda-Arara, B. 2014. Genetic diversity and gene flow in *Clarias gariepinus* from Lakes Victoria and Kanyaboli, Kenya. African Journal of Aquatic Science, 39, 287–293. 10.2989/16085914.2014.933734

Barasa, J. E., Mdyogolo, S., Abila, R., Grobler, J. P., Skilton, R. A., Bindeman, H., Njahira, M. N., Chemoiwa, E. J., Dangasuk, O. G., Kaunda-Arara, B. & Verheyen, E. 2017. Genetic diversity and population structure of the African catfish, Clarias gariepinus (Burchell, 1822) in Kenya: implication for conservation and aquaculture. Belgian Journal of Zoology, 147, 105-127. 10.26496/bjz.2017.9

Bayona-Vásquez, N. J., Glenn, T. C., Kieran, T. J., Pierson, T. W., Hoffberg, S. L., Scott, P. A., Bentley, K. E., Finger, J. W., Louha, S. & Troendle, N. 2019. Adapterama III: Quadruple-indexed, double/triple-enzyme RADseq libraries (2RAD/3RAD). PeerJ, 7, e7724. 10.7717/peerj.7724

Behmene, I. E., Bachir Bouiadjra, B., Homrani, A., Daoudi, M., Sánchez-Vázquez, F. J., López-Lopez, A., Asensio-Pérez, A. I. & Galián, J. 2022. Morphometric and genetic diversity of an African catfish (*Clarias gariepinus*) population from Southeast Algeria. African Journal of Ecology, 60, 1287–1292. 10.1111/aje.13055

Bernatchez, L., Wellenreuther, M., Araneda, C., Ashton, D. T., Barth, J. M. I., Beacham, T. D., Maes, G. E., Martinsohn, J. T., Miller, K. M., Naish, K. A., Ovenden, J. R., Primmer, C. R., Young Suk, H., Therkildsen, N. O. & Withler, R. E. 2017. Harnessing the power of genomics to secure the future of seafood. Trends in Ecology & Evolution, 32, 665–680. 10.1016/j.tree.2017.06.010

Bian, C., Huang, Y., Li, J., You, X., Yi, Y., Ge, W. & Shi, Q. 2019. Divergence, evolution and adaptation in ray-finned fish genomes. Science China Life Sciences, 62, 1003–1018. 10.1007/s11427-018-9499-5

Chandra Segaran, T., Azra, M. N., Piah, R. M., Lananan, F., Téllez-Isaías, G., Gao, H., Torsabo, D., Kari, Z. A. & Noordin, N. M. 2023. Catfishes: A global review of the literature. Heliyon, 9, e20081. 10.1016/j.heliyon.2023.e20081

Coleman, R. A., Gauffre, B., Pavlova, A., Beheregaray, L. B., Kearns, J., Lyon, J., Sasaki, M., Leblois, R., Sgro, C. & Sunnucks, P. 2018. Artificial barriers prevent genetic recovery of small isolated populations of a low-mobility freshwater fish. Heredity, 120, 515–532. 10.1038/s41437-017-0008-3

Crollius, H. R. & Weissenbach, J. 2005. Fish genomics and biology. Genome Research, 15, 1675–1682. 10.1101/gr.3735805

Dumith, M. T. & Santos, A. F. G. N. D. 2022. Ecological Impacts of the African catfish *Clarias gariepinus* at Environmental Protection Area in southeastern Brazil. Research Square Platform llc. 10.21203/rs.3.rs-1132915/v1

Duncan, N. J., Sonesson, A. K. & Chavanne, H. 2013. 2 - Principles of finfish broodstock management in aquaculture: control of reproduction and genetic improvement. *In:* Allan, G. & Burnell, G. (eds.) Advances in Aquaculture Hatchery Technology. Woodhead Publishing. 10.1533/9780857097460.1.23

Emmanuel, E., Ekpe, A. & Okon, A. 2021. Threatened and endangered fish species in Nigeria, a menace to biodiversity – A review. African Journal of Education, Science and Technology, 3, 12–26. 10.2022/ajest.v3i1.665

Erkinaro, J., Niemelä, E., Vähä, J.-P., Primmer, C. R., Brørs, S. & Hassinen, E. 2010. Distribution and biological characteristics of escaped farmed salmon in a major subarctic wild salmon river: implications for monitoring. Canadian Journal of Fisheries and Aquatic Sciences, 67, 130–142. 10.1139/f09-173

Essien, E., Jesse, E. & Igbokwe, J. 2019. Assessment of water level in Dadin Kowa Dam reservoir in Gombe State Nigeria using geospatial techniques. International Journal of Environment and Geoinformatics, 6, 115–130. 10.30897/ijegeo.487885

Excoffier, L. & Lischer, H. E. L. 2010. Arlequin suite ver 3.5: a new series of programs to perform population genetics analyses under Linux and Windows. Molecular Ecology Resources, 10, 564–567. 10.1111/j.1755-0998.2010.02847.x

Eyo, V. O., Eriegha, O. J. & Eze, F. 2021. Reproductive performance of hatchery-bred, wild-caught broodstock, and their outbreed of the African catfish Clarias gariepinus (Burchell, 1822). Indonesian Aquaculture Journal, 15, 59-65. 10.15578/iaj.15.2.2020.59-65

Eze, F., Haruna, M. Y. & Victory, A. O. 2022. A comparative study on the gonadosomatic index and milt volume of four populations of Clarias gariepinus (Burchell, 1822) broodstock strains from North-East Nigeria. Asian Journal of Fisheries and Aquatic Research, 17, 8-19. 10.9734/ajfar/2022/v17i530415

Ezilrani, P. & Christopher, J. G. 2015. Genetic variation and differentiation in African catfish, Clarias gariepinus, assessed by heterologous microsatellite DNA. Indian Journal of Biotechnology, 14, 388–393.

FAO 2022. The State of World Fisheries and Aquaculture 2022. Towards Blue Transformation. Food & Agriculture Organization of the United Nations, Rome.

Fazzi-Gomes, P. F., Aguiar, J. d. P., Marques, D., Fonseca Cabral, G., Moreira, F. C., Rodrigues, M. D. N., Silva, C. S., Hamoy, I. & Santos, S. 2021. Novel microsatellite markers used for determining genetic diversity and tracing of wild and farmed populations of the Amazonian giant fish *Arapaima gigas*. Genes, 12, 1324. 10.3390/genes12091324

Fernández, J., Toro, M. Á., Sonesson, A. K. & Villanueva, B. 2014. Optimizing the creation of base populations for aquaculture breeding programs using phenotypic and genomic data and its consequences on genetic progress. Frontiers in Genetics, 5. 10.3389/fgene.2014.00414

Folorunso, E. A., Rahman, M. A., Sarfo, I., Darko, G. & Olowe, O. S. 2021. Catfish farming: a sustainability study at Eriwe fish farming village in southwest Nigeria. Aquaculture International, 29, 827–843. 10.1007/s10499-021-00662-0

Goudet, J. 2005. hierfstat, a package for r to compute and test hierarchical F-statistics. Molecular Ecology Notes, 5, 184–186. 10.1111/j.1471-8286.2004.00828.x

Hassan, A., Adewumi, M. O., Ayeni, M. D. & Falola, A. 2015. An assessment of the irrigation scheme on registered rice farmers of the Upper Benue rice basin development in Dadin Kowa, Gombe State Nigeria. Journal of Multidisciplinary Studies, 4. 10.7828/jmds.v4i1.843

Hecht, T. 2013. A review of on-farm feed management practices for North African catfish (*Clarias gariepinus*) in sub-Saharan Africa. On-farm Feeding and Feed Management in Aquaculture. Rome: FAO Fisheries and Aquaculture, Rome.

Holčík, J. 1991. Fish introductions in Europe with particular reference to its central and eastern part. Canadian Journal of Fisheries and Aquatic Sciences, 48, 13–23. 10.1139/f91-300

Hughes, L. C., Ortí, G., Huang, Y., Sun, Y., Baldwin, C. C., Thompson, A. W., Arcila, D., Betancur-R., R., Li, C., Becker, L., Bellora, N., Zhao, X., Li, X., Wang, M., Fang, C., Xie, B., Zhou, Z., Huang, H., Chen, S., Venkatesh, B. & Shi, Q. 2018. Comprehensive phylogeny of ray-finned fishes (Actinopterygii) based on transcriptomic and genomic data. Proceedings of the National Academy of Sciences, 115, 6249–6254. 10.1073/pnas.1719358115

Huisman, E. & Richter, C. 1987. Reproduction, growth, health control and aquacultural potential of the African catfish, Clarias gariepinus (Burchell 1822). *Aquaculture*, 63, 1-14. 10.1016/0044-8486(87)90057-3

Hvilsom, C., Segelbacher, G., Ekblom, R., Fischer, M. C., Laikre, L., Leus, K., ÓBrien, D., Shaw, R. & Sork, V. 2022. Selecting species and populations for monitoring of genetic diversity, Gland, Switzerland: IUCN, International Union for Conservation of Nature.

Ibiwoye, Y. E. 2018. Assessment of broodstock management practices in Nigeria. United Nations University, Iceland.

Iyiola, O. A., Nneji, L. M., Mustapha, M. K., Nzeh, C. G., Oladipo, S. O., Nneji, I. C., Okeyoyin, A. O., Nwani, C. D., Ugwumba, O. A. & Ugwumba, A. A. 2018. DNA barcoding of economically important freshwater fish species from north-central Nigeria uncovers cryptic diversity. Ecology and Evolution, 8, 6932–6951. 10.1002/ece3.4210

Jombart, T. 2008. Adegenet: A R package for the multivariate analysis of genetic markers. Bioinformatics, 24, 1403–1405. 10.1093/bioinformatics/btn129

Jombart, T., Kamvar, Z. N., Collins, C., Luštrik, R., Beugin, M.-P., Knaus, B. J., Solymos, P., Mikryukov, V., Schliep, K. & Maié, T. 2023. Adegenet: Exploratory analysis of genetic and genomic data v2.1.10.

Kemp, P. S., Subbiah, G., Barnes, R., Boerder, K., O’Leary, B. C., Stewart, B. D. & Williams, C. 2023. The future of marine fisheries management and conservation in the United Kingdom: Lessons learnt from over 100 years of biased policy. Marine Policy, 147, 105075. 10.1016/j.marpol.2022.105075

Kenchington, E. 2003. The Effects of Fishing on Species and Genetic Diversity. *In:* Sinclair, M. & Valdimarson, G. (eds.) Responsible Fisheries in the Marine Ecosystem. Wallingford, Oxon, UK: CAB International. 10.1079/9780851996332.0235

Kolde, R. 2019. pheatmap: Pretty heatmaps. R package version 1.0.12.

Konings, A., Freyhof, J., FishBase team RMCA & Geelhand, D. 2019. *Clarias gariepinus* (amended version of 2018 assessment). The IUCN Red List of Threatened Species 2019: e.T166023A155051767. Available on: 10.2305/IUCN.UK.2018-2.RLTS.T166023A155051767.en.

Kundu, S., De Alwis, P. S., Binarao, J. D., Lee, S.-R., Kim, A. R., Gietbong, F. Z., Yi, M. & Kim, H.-W. 2023. Mitochondrial DNA corroborates the genetic variability of *Clarias* catfishes (Siluriformes, Clariidae) from Cameroon. Life, 13, 1068. 10.3390/life13051068

Lal, K., Singh, R., Mohindra, V., Singh, B. & Ponniah, A. 2003. Genetic make up of exotic catfish *Clarias gariepinus* in India. Asian Fisheries Science, 16, 229–234. https://hdl.handle.net/20.500.12348/2216

Larsson, A. 2014. AliView: a fast and lightweight alignment viewer and editor for large datasets. Bioinformatics, 30, 3276–3278. 10.1093/bioinformatics/btu531

Li, H. & Durbin, R. 2009. Fast and accurate short read alignment with Burrows-Wheeler transform. Bioinformatics, 25, 1754–60. 10.1093/bioinformatics/btp324

MacKenzie, S. & Jentoft, S. 2016. 11 - Future perspective. *In:* Mackenzie, S. & Jentoft, S. (eds.) Genomics in Aquaculture. San Diego: Academic Press. 10.1016/B978-0-12-801418-9.00011-1

Mahlum, S., Vollset, K. W., Barlaup, B. T., Skoglund, H. & Velle, G. 2021. Salmon on the lam: Drivers of escaped farmed fish abundance in rivers. Journal of Applied Ecology, 58, 550–561. 10.1111/1365-2664.13804

Martinez, A. S., Willoughby, J. R. & Christie, M. R. 2018. Genetic diversity in fishes is influenced by habitat type and life-history variation. Ecology and Evolution, 8, 12022–12031. 10.1002/ece3.4661

Mekuleyi, G. O., Awe, F. A., Whenu, O. O., Adeboyejo, O. A., Hungbo, J. J., Akinyemi, A. A. & Olanloye, O. A. 2022. Isolation and identification of bacteria found in the milt of cultured *Clarias gariepinus*. Asian Journal of Fisheries and Aquatic Research, 4, 8–19. 10.9734/ajfar/2022/v17i530415

Meulenbroek, P., Curto, M., Priglinger, P., Pinter, K., Shumka, S., Graf, W., Schiemer, F. & Meimberg, H. 2024. Small-scale metapopulation structure of a limnophilic fish species in a natural river system investigated using microsatellite genotyping by amplicon sequencing (SSR-GBAS). BMC Ecology and Evolution, 24, 1. 10.1186/s12862-023-02192-0

Mramba, R. P. & Kahindi, E. J. 2023. The status and challenges of aquaculture development in Dodoma, a semi-arid region in Tanzania. Aquaculture International, 31, 1551–1568. 10.1007/s10499-022-01041-z

Ng, P. C. & Kirkness, E. F. 2010. Whole genome sequencing. Methods in Molecular Biology, 628, 215–26. 10.1007/978-1-60327-367-1_12

Nwani, C. D., Becker, S., Braid, H. E., Ude, E. F., Okogwu, O. I. & Hanner, R. 2011. DNA barcoding discriminates freshwater fishes from southeastern Nigeria and provides river system-level phylogeographic resolution within some species. Mitochondrial DNA, 22 Suppl 1, 43–51. 10.3109/19401736.2010.536537

Nyunja, C., Maina, J., Amimo, J., Kibegwa, F., Harper, D. & Jung’a, J. 2017. Stock structure delineation of the African Catfish (*Clarias gariepinus*) in selected populations in Kenya using mitochondrial DNA (Dloop) variability. Journal of Aquaculture Research & Development, 8, 1–6. 10.4172/2155-9546.1000485

Ola-Oladimeji, F. A., Oso, J. A., Oladimeji, T. E., Idowu, E. O., Adeleke, K. & Urihe, F. O. 2017. Phenotypic diversities of four populations of *Clarias gariepinus* (Siluriformes, Clariidae) obtained from Ogun and Ondo State waterbodies in south-western Nigeria. Vestnik Zoologii, 51, 285–294. 10.1515/vzoo-2017-0034

Olopade, O., Taıwo, İ. & Dıenye, H. 2017. Management of overfishing in the inland capture fisheries in Nigeria. Journal of Limnology and Freshwater Fisheries Research, 3, 189–194. 10.17216/limnofish.335549

Omitoyin, B. O. 2007. Introduction to Fish Farming in Nigeria, Ibadan University Press.

Parvez, I., Rumi, R. A., Ray, P. R., Hassan, M. M., Sultana, S., Pervin, R., Suwanno, S. & Pradit, S. 2022. Invasion of African *Clarias gariepinus* drives genetic erosion of the indigenous *C. batrachus* in Bangladesh. Biology, 11, 1–12. 10.3390/biology11020252

Peterson, B. K., Weber, J. N., Kay, E. H., Fisher, H. S. & Hoekstra, H. E. 2012. Double digest RADseq: an inexpensive method for de novo SNP discovery and genotyping in model and non-model species. PloS one, 7, e37135. 10.1371/journal.pone.0037135

Petit-Marty, N., Liu, M., Tan, I. Z., Chung, A., Terrasa, B., Guijarro, B., Ordines, F., Ramírez-Amaro, S., Massutí, E. & Schunter, C. 2022. Declining population sizes and loss of genetic diversity in commercial fishes: A simple method for a first diagnostic. Frontiers in Marine Science, 9. 10.3389/fmars.2022.872537

Popoola, O. M. 2022. Genetic differentiation and molecular phylogenetics of North African catfish from three distinct waterbodies. Croatian Journal of Fisheries, 80, 123–132. 10.2478/cjf-2022-0013

Popoola, O. M., Fasakin, A. E. & Awopetu, I. J. 2014. Genetic variability in cultured and wild population of *Clarias gariepinus* using Sodium Dodecyl Sulfate-Polyacrylamide Gel Electrophoresis (SDS-PAGE). Biotechnology Journal International, 72, 5–11. 10.14798/72.1.712

R Core Team 2018. R: A language and environment for statistical computing. R Foundation for Statistical Computing, Vienna, Austria. Available online at: https://www.R-project.org/. Vienna, Austria.

Rivera-Colón, A. G. & Catchen, J. 2022. Population Genomics Analysis with RAD, Reprised: Stacks 2. *In:* Verde, C. & Giordano, D. (eds.) Marine Genomics: Methods and Protocols. New York, NY: Springer US. 10.1007/978-1-0716-2313-8_7

Roodt-Wilding, R., Swart, B. L. & Impson, N. D. 2010. Genetically distinct Dutch-domesticated *Clarias gariepinus* used in aquaculture in southern Africa. African Journal of Aquatic Science, 35, 241–249. 10.2989/16085914.2010.538507

Rozas, J., Ferrer-Mata, A., Sánchez-DelBarrio, J. C., Guirao-Rico, S., Librado, P., Ramos-Onsins, S. E. & Sánchez-Gracia, A. 2017. DnaSP 6: DNA sequence polymorphism analysis of large data sets. Molecular Biology and Evolution, 34, 3299–3302. 10.1093/molbev/msx248

Sanda, M. K., Metcalfe, N. B. & Mable, B. K. 2024. The potential impact of aquaculture on the genetic diversity and conservation of wild fish in sub-Saharan Africa. Aquatic Conservation: Marine and Freshwater Ecosystems, 34, e4105. 10.1002/aqc.4105

Šegvić-Bubić, T., Grubišić, L., Trumbić, Ž., Stanić, R., Ljubković, J., Maršić-Lučić, J. & Katavić, I. 2016. Genetic characterization of wild and farmed European seabass in the Adriatic sea: assessment of farmed escapees using a Bayesian approach. ICES Journal of Marine Science, 74, 369–378. 10.1093/icesjms/fsw155

Sonet, G., Snoeks, J., Nagy, Z. T., Vreven, E., Boden, G., Breman, F. C., Decru, E., Hanssens, M., Ibala Zamba, A., Jordaens, K., Mamonekene, V., Musschoot, T., Van Houdt, J., Van Steenberge, M., Lunkayilakio Wamuini, S. & Verheyen, E. 2019. DNA barcoding fishes from the Congo and the Lower Guinean provinces: Assembling a reference library for poorly inventoried fauna. Molecular Ecology Resources, 19, 728–743. 10.1111/1755-0998.12983

Suleiman, S. A. B., Diyaware, M. Y., Mohammed, Z. B. & Aliyu, M. 2020. Genetic characterization of farmed and wild populations of African catfish (*Clarias gariepinus* Burchell, 1822) using the random amplified polymorphic marker Journal of Agricultural Sciences, Belgrade, 65, 375–389. 10.2298/jas2004375s

Tadmor-Levi, R., Feldstein-Farkash, T., Milstein, D., Golani, D., Leader, N., Goren, M. & David, L. 2023. Revisiting the species list of freshwater fish in Israel based on DNA barcoding. Ecology and Evolution, 13, e10812. 10.1002/ece3.10812

Truter, M., Hadfield, K. A. & Smit, N. J. 2023. Chapter Two - Review of the metazoan parasites of the economically and ecologically important African sharptooth catfish Clarias gariepinus in Africa: Current status and novel records. *In:* Rollinson, D. & Stothard, R. (eds.) Advances in Parasitology. Academic Press. 10.1016/bs.apar.2022.11.001

Turan, F. & Turan, C. 2016. Natural and non-natural distribution of African catfish Clarias gariepinus (Burchell, 1822) in Turkey. Journal of Limnology and Freshwater Fisheries Research, 2, 173-177. 10.17216/limnofish.280413

Wachirachaikarn, A. & Na-Nakorn, U. 2019. Genetic diversity of the North African catfish, Clarias gariepinus (Burchell, 1822) hatchery stocks in Thailand. Science Asia, 45, 301-308. 10.2306/scienceasia1513-1874.2019.45.301

Weir, B. S. & Cockerham, C. C. 1984. Estimating F-statistics for the analysis of population structure. Evolution, 38, 1358–1370. 10.1111/j.1558-5646.1984.tb05657.x

Wickham, H. 2016. ggplot2: Elegant Graphics for Data Analysis, Springer-Verlag New York.

Yan, H. F., Kyne, P. M., Jabado, R. W., Leeney, R. H., Davidson, L. N. K., Derrick, D. H., Finucci, B., Freckleton, R. P., Fordham, S. V. & Dulvy, N. K. 2021. Overfishing and habitat loss drive range contraction of iconic marine fishes to near extinction. Science Advances, 7, eabb6026. doi:10.1126/sciadv.abb6026

Zeinab, Z., Shabany, A. & Kolangi-Miandare, H. 2014. Comparison of genetic variation of wild and farmed Bream (*Abramis brama orientalis*; berg, 1905) using microsatellite markers. Molecular Biology Research Communications, 3, 187–195.

