## Supplementary material for "Genetic diversity, population structure and differentiation of farmed and wild African catfish (*Clarias gariepinus*) in Nigeria": https://github.com/mksanda/Clarias_gariepinus_3RAD

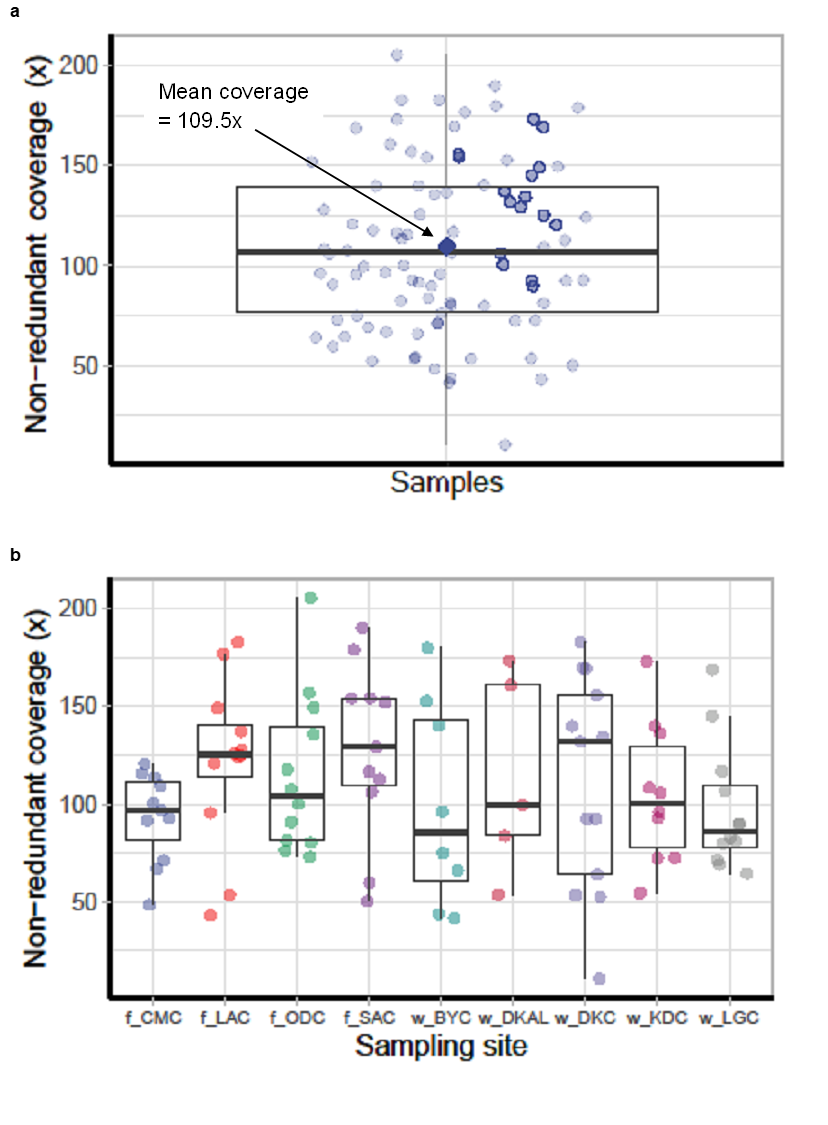

**Supplementary Figure 1.** Boxplot illustrating the distribution of non-redundant read coverage indicating the number of reads mapped to specific genomic regions or features for each population for individual samples with mean coverage (a), and distribution of reads across sampling sites (b). The boxplot provides a visual summary of the read distribution, with the box indicating the interquartile range (IQR) and the median read coverage, and the whiskers extending to the minimum and maximum values or to a specified range. The prefixes f and w, on the x-axis indicate farmed and wild samples, respectively.

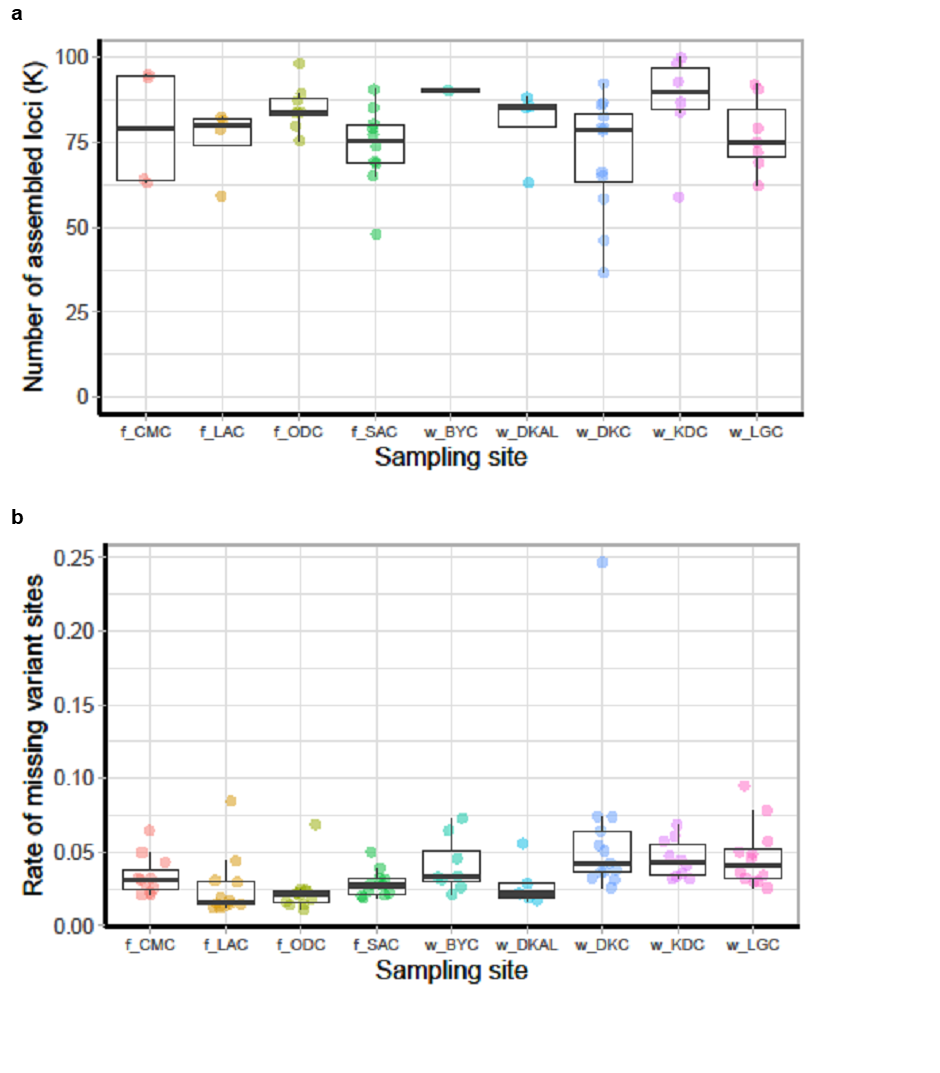

**Supplementary Figure 2.** Boxplot illustrating the distribution of assembled loci across different sampling locations (a) and the rate of missing variant sites (b) for the 3RAD sequencing data. The y-axis indicates the number of loci in thousands and the x-axis represents the sampling locations. Each point on the plot represents a sample within a location. The prefixes f and w, on the x-axis indicate farmed and wild samples, respectively.

**Supplementary Table 1. Global distribution of haplotypes based on 100% match from BOLD Systems search.**

| **Haplotype** | **Country and reference** |
| --- | --- |
| 5 | Nigeria (n = 2): MG824580, MG824583 (Iyiola et al., 2018), BAFEN141-10 (Nwani et al., 2011b); Algeria (n = 4): ON643478, ON643477, ON643476, ON643475 (Behmene et al., 2022); Egypt (n = 1): MK335911 (unpublished) |
| 7 | Nigeria (n = 2): MG824581 (Iyiola et al., 2018), BAFEN140-10 (Nwani et al., 2011b) |
| 8 | Israel (n = 11): FWISR057-21, FWISR056-21, FWISR055-21, FWISR054-21, FWISR053-21, FWISR052-21, FWISR046-21, LKCOX059-19, LKCOX058-19, LKCOX057-19, LKCOX056-19 (Tadmor-Levi et al., 2023); Thailand (n = 3): JF292311, JF292314, MT571809 (Wong et al., 2011); Bangladesh (n = 1): MG988400 (unpublished); Egypt (n = 2): MK335909, MK335910, (Unpublished); Nigeria (n = 12): BAFEN129-10, BAFEN130-10, BAFEN131-10, BAFEN134-10, BAFEN137-10, BAFEN144-10, BAFEN145-10, BAFEN146-10, BAFEN147-10, BAFEN148-10, BAFEN150-10, BAFEN151-10 (Nwani et al., 2011b); Syria (n = 1): FFMBH2002-14 (Geiger et al., 2014); India (n = 1): FNWG200-16 (Patil et al., 2018) |
| 9 | Israe (n = 7): FWISR057-21, FWISR056-21, FWISR055-21, FWISR054-21, FWISR053-21, FWISR052-21, FWISR046-21 (Tadmor-Levi et al., 2023); Thailand (n = 1): JF292314 (Wong et al., 2011); Bangladesh (n = 1): MG988400 (unpublished); Egypt (n = 1): MK335909, MK335910 (Unpublished) |
| 10 | DR Congo (n = 5): BCOVR501-17 (Sonet et al., 2019), DCF305-15, DCF751-15, DCF752-1, DCF602-15 (unpublished); Brazil (n = 6): FUPR532-09, LBPV-31863, LBPV-31864, LBPV-31865, LBPV-31866, FUPR536-09 (Pereira et al., 2013); |
| 11 | DR Congo (n = 1) BIN ID: AAB2256 (Sonet et al., 2019) |

**Supplementary Table 2. Analysis of Molecular Variance (AMOVA) of mtDNA COI sequences in farmed and wild *C. gariepinus* populations from northeastern and southwestern Nigeria including albino as wild samples**

| **Source of Variation** | **d.f.** | **SS** | **Var** | **% Var** |
| --- | --- | --- | --- | --- |
| Among groups | 2 | 8.992 | -0.046 Va | -11.17 |
| Among populations within groups | 8 | 18.083 | 0.169 Vb | 41.35 |
| Within populations | 200 | 57.118 | 0.286 Vc | 69.82 |
| Total | 210 | 84.123 | 0.409 | 100 |

Notes: SS, sum of squares; Var, variance component; Va, variance components among populations; Vb, variance components among populations within groups; Vc, variance components within populations

**Supplementary Table 3. Analysis of Molecular Variance (AMOVA) of mtDNA COI sequences in farmed and wild *C. gariepinus* populations from northeastern and southwestern Nigeria without including the albino sample**

| **Source of Variation** | **d.f.** | **SS** | **Var** | **% Var** |
| --- | --- | --- | --- | --- |
| Among groups | 2 | 9.995 | -0.039 Va | -9.35 |
| Among populations within groups | 7 | 17.278 | 0.174 Vb | 42.16 |
| Within populations | 192 | 53.302 | 0.278 Vc | 67.19 |
| Total | 201 | 80.575 | 0.413 | 100 |

Notes: SS, sum of squares; Var, variance component; Va, variance components among populations; Vb, variance. components among populations within groups; Vc, variance components within populations.
